# CellWHISPER disentangles direct cell–cell communication from structural proximity

**DOI:** 10.64898/2026.01.07.697982

**Authors:** Anurendra Kumar, Felix Rivera, Bhavay Aggarwal, Nicholas Zhang, Ahmet Coskun, Saurabh Sinha

## Abstract

Spatial transcriptomics makes it possible to infer direct, contact-dependent cellular communication, but existing tools are prone to high error rates due to spatial structure in cell types and gene expression. We present CellWHISPER, a framework for inferring contact-mediated communication, including gap junctions and ligand-receptor signaling, from spatial transcriptomics data. CellWHISPER controls errors with a statistical procedure that accounts for confounding effects of cell-type-specific expression and spatial organization. Closed-form significance scores enable analysis of large data. A latent variable model distills recurrent patterns into maps of preferences among cell types and signaling genes. We demonstrate CellWHISPER on two technologies (Stereo-seq, Xenium), two species and three tissues. In mouse brain it yields a comprehensive connexin-interaction map. Immunostaining confirmed CellWHISPER-predicted Connexin 43-mediated interfaces among astrocytes, microglia and endothelial cells, which existing approaches fail to prioritize. In human colon and skin, tissues with strong spatial gradients, CellWHISPER recovers communication mechanisms both along and independent of the gradient.

## INTRODUCTION

Direct contact-mediated cell signaling shapes tissue homeostasis and disease, yet existing computational tools focus almost exclusively on paracrine mechanisms. Rapid proliferation of single-cell transcriptomics technologies has led to several methods for discovering cell-cell communication (CCC) mechanisms^1–4^. These tools, such as CellChat^5^, CellPhoneDB^6^ and NicheNet^7^, typically reveal communication between specific cell types via curated ligand-receptor (LR) pairs. Broadly, CCC can be categorized into paracrine, autocrine, juxtacrine, and endocrine signaling^1^. Existing CCC methods focus on paracrine and autocrine signaling mediated by LR interactions, whereas direct juxtacrine signaling, particularly via gap junction communication (GJC), remains largely unexplored. Rapidly emerging spatial transcriptomics (ST) technologies including Stereoseq, VisiumHD, Nanostring and Xenium, provide a transformative opportunity to bridge this gap by combining gene expression with spatial proximity, enabling investigation of direct (contact-mediated) communication between cells. However, computational inference of direct CCC poses distinct statistical challenges: spatial organization of cell-types and heterogeneous gene expression act as strong confounders that can inflate apparent evidence for signaling if not explicitly controlled (**Fig. 1a**).

**Figure 1.**
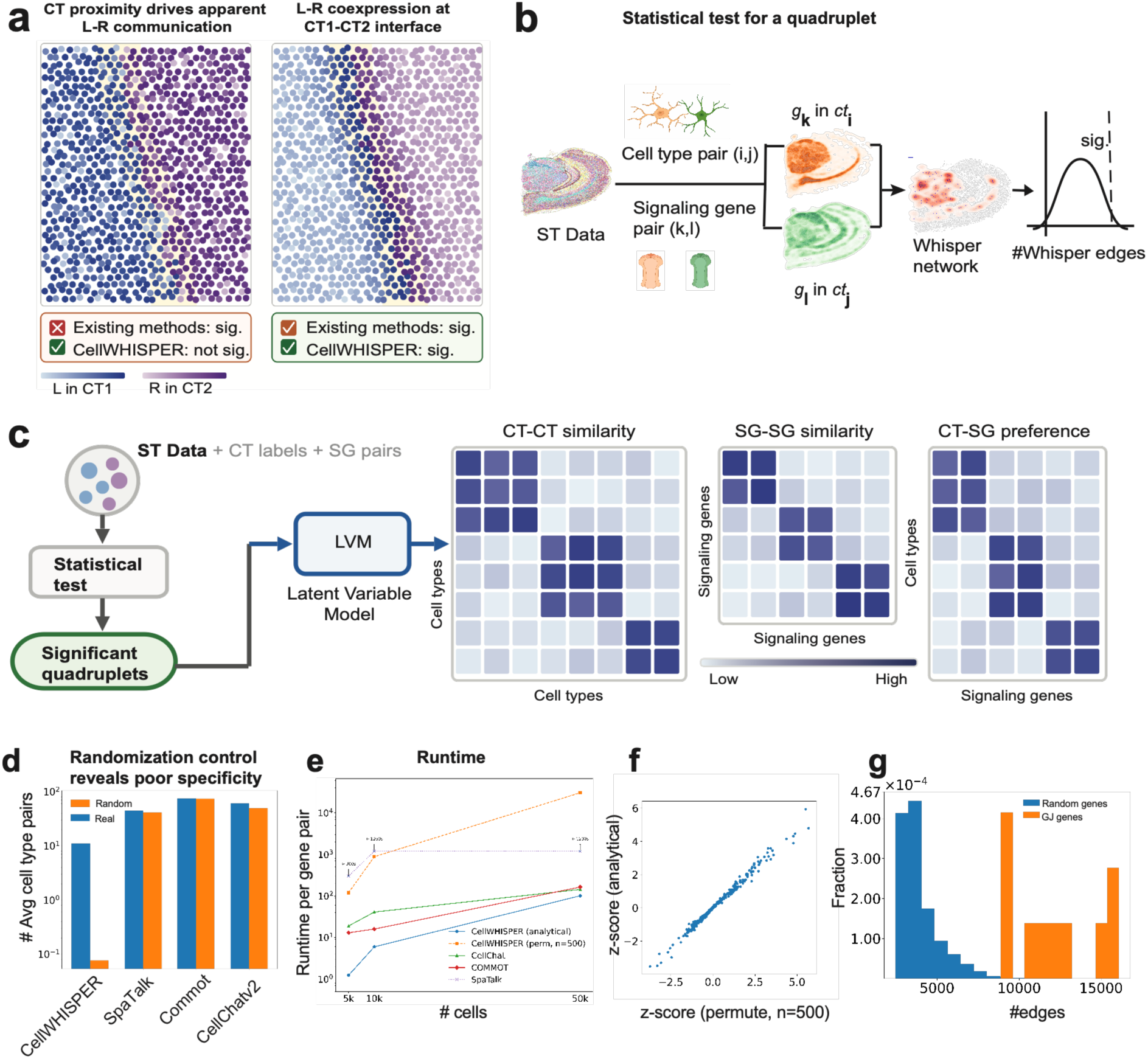
Overview of CellWHISPER. (**a**) **Cell-type confounding in ligand–receptor communication inference.** When ligand L is expressed broadly across cell type CT1 and receptor R across CT2, spatial proximity between the two cell types produces apparent L–R co-expression at their interface (left; yellow zone) even without specific L–R signaling. When the same number of high-expressing cells are enriched specifically at the CT1–CT2 interface, the co-expression reflects genuine signaling (right; yellow zone). Both scenarios appear identical to existing methods, which report significant communication in both cases. CellWHISPER evaluates co-expression against a null that randomizes cell identities within each cell type, preserving cell-type-specific expression distributions and spatial organization, and correctly distinguishes the two. (**b**) **Statistical testing for a CT-SG quadruplet**: For a given cell type pair (*ct_i_*, *ct*_j_) and signaling gene pair (*g_k_*, *g_l_*), CellWHISPER constructs a “whisper network”, where each edge represents a pair of adjacent cells of types *ct_i_*, *ct*_j_ expressing genes *g_k_*, *g_l_* respectively, and the count of such “whisper edges” serves as the test statistic. Under the null model, cell locations are randomized within each cell type, generating a background distribution of edge counts, which is used to assess statistical significance. (**c**) **CellWHISPER framework**: CellWHISPER takes single-cell resolution ST data as input, along with cell type (CT) labels and signaling gene pairs, which may be ligand-receptor (LR) gene pairs or gap junction (GJ) gene pairs. It performs a statistical test (panel b) to compute a z-score for each cell type pair and signaling gene pair (a CT-SG quadruplet), reporting all quadruplets with significant z-scores. A latent variable model can then analyze the full complement of quadruplet-level results to infer higher-order communication patterns such as CT pairs that are similar in their SG usage (CT-CT similarity matrix), SG pairs that mediate communication between similar cell types (SG-SG similarity matrix), and quantify whether a certain cell types tends to communicate using a specific SG (CT-SG preference matrix). CellWHISPER can also be applied across conditions for differential analysis, identifying CT-SG quadruplets specific to one condition versus the other. (**d**) **Randomized control reveals poor specificity**: Shown is the average number of communicating cell-type pairs inferred per ligand–receptor by four different methods, on real versus randomized data (cell identities permuted within each cell type, preserving expression distributions and cell-type spatial organization while disrupting cross-type proximity). Competing methods show limited separation between real and randomized data, whereas CellWHISPER yields substantially fewer calls under randomization, indicating significantly improved specificity for direct CCC. We summarize this as an empirical FPR, defined as the number of significant CT–SG quadruplets detected on randomized data divided by the number detected on real data under fixed thresholds (CellWHISPER: <5%). **Benchmarking was performed on a downsampled hippocampal subset (∼5,000 cells) due to computational constraints of competing methods** (Methods, Supp. Fig. 5). (**e**) **Runtime and scalability.** Wall-clock time to evaluate all cell-type pairs for a single signaling-gene pair on datasets of increasing size (∼5,000; ∼10,000; and ∼50,000 cells), comparing CellWHISPER with the analytical null versus permutation testing (n = 500), and three competing tools (CellChat, COMMOT, SpaTalk) run with recommended settings. CellWHISPER’s analytical null achieves ∼100× speedup over permutation on ∼5,000 cells and substantially larger speedups at ∼10,000 and ∼50,000 cells, enabling practical screening on full-resolution tissues. SpaTalk did not complete within the allotted runtime at one or more dataset sizes (shown as timeouts). **(f) Consistency between analytical and permutation-based significance**. Z-scores computed using the analytical null closely match those from permutation testing across CT–SG tests, confirming that the analytical formulation reproduces the permutation-based null while avoiding explicit shuffling. (**g**) **Whisper network specificity for connexins:** For each homotypic connexin gene, an aggregated whisper network was constructed as the union of whisper networks across all cell-type pairs in the Stereo-seq mouse brain dataset. Edge counts for connexin networks were significantly higher than those for networks constructed from randomly selected genes (n = 5000) (Mann–Whitney U test p = 4.6 × 10e-8). All connexins yielded ≥ 8,878 edges, whereas only 0.4% of random genes reached this threshold.

Direct cell-cell signaling operates primarily via two mechanisms: GJC^8^ and LR interactions. Gap junctions are intercellular channels (**Supp. Fig. 7**) that enable direct cell-cell transfer of ions, metabolites, second messengers, and regulatory molecules^9^. GJC-mediated inter-cellular networks play vital roles in normal physiology^10–12^ and are implicated in diverse pathological conditions^13–15^. Gap junction channels are formed by connexin proteins, assembled as hexamers of homotypic or heterotypic connexins. Over 20 connexins have been identified in mammals, and the existence of a compatibility “code” governing gap junction formation is an important open problem^16,17^. Similarly, direct LR pathways, such as Notch-Delta^18^ and Eph-Ephrin^19^ regulate differentiation, embryogenesis, and immune cell communication^20^, motivating the search for additional instances of direct LR signaling in varying contexts^21^.

Numerous tools exist for CCC inference from single-cell transcriptomics, including CellChat, CellPhoneDB, NicheNet, and LIANA+, among others. However, these tools infer paracrine signaling from co-expression patterns alone and are fundamentally unsuited for contact-mediated communication, which requires physical proximity between cells. Very few tools, specifically CellChatv2^22^, SpaTalk^23^, and COMMOT^24^, exploit inter-cellular proximity information available in ST data, a key requirement for inferring *contact-dependent* communication. We found these ST-based inference tools to exhibit surprisingly high false positive errors, motivating this study. Specifically, we performed a randomized control by permuting cell locations within each cell type, preserving cell-type-specific spatial organization and LR expression while destroying spatial proximity between ligand- and receptor-expressing cells. In this control, evidence for true direct CCC should be eliminated. Strikingly, the above-mentioned ST-based CCC inference tools reported comparable numbers of direct CCC mechanisms in randomized and original datasets, indicating insufficient statistical calibration for spatially constrained inference. This finding is consistent with published studies debating the error rates of existing tools ^3,25^ and suggestions of inflated false positives^1,4,26^. It motivated us to devise a statistical procedure for CCC inference that explicitly accounts for gene-expression heterogeneity, spatial organization, and statistical dependencies induced by shared cellular neighborhoods.

We present CellWHISPER (<u>W</u>orkflow for <u>HI</u>gh-precision <u>S</u>patial <u>P</u>roximity-mediated c<u>E</u>ll-cell inte<u>R</u>actions), a toolkit to infer direct CCC from single-cell ST data. CellWHISPER makes four core contributions: (i) a stringent benchmarking strategy that isolates contact-dependent signal from confounding effects of cell-type, revealing that state-of-the-art ST-based tools fail to control false positives, (ii) a proximity-constrained statistical test with a confounder-aware null that improves specificity for direct CCC, (iii) an analytically derived null (closed-form mean/variance of the CCC-measuring statistic) that avoids explicit permutations and enables scalable inference across large tissues and signaling compendia, and (iv) a latent variable model that analyzes the full tensor of tested CCC mechanisms to capture its higher-order interpretable structures without first collapsing the tensor into information-losing matrices (as is currently done). Applying CellWHISPER to mouse brain datasets from STEREO-seq^27^ and Xenium recovers known and predicts and novel GJC, reporting the first comprehensive brain “connexin code”^16,28^. Immunostaining furnished supporting evidence for Cx43-mediated interfaces among astrocytes, microglia and endothelial cells, which were among the highest-ranked predictions; in contrast, existing approaches fail to prioritize these confirmed interactions. LR analysis recapitulates established CCC programs such as mossy fiber pathway while a comparison of an Alzheimer’s disease (AD) model with wild-type brain highlights AD-associated microglial GJC. Application to human colon and skin, tissues with strong spatial gradients, shows that CellWHISPER recovers contact circuits along and independent of the gradient without being confounded by zonal co-expression. In summary, CellWHISPER enables rigorous, scalable inference of direct inter-cellular communication, providing a brain-wide connexin-interaction map.

## RESULTS

### Overview of CellWHISPER

CellWHISPER infers direct cell-cell communication (CCC) involving a “quadruplet” of two cell types and two signaling genes. Given ST data and cell-type labels, it tests whether a cell-type pair shows evidence of direct communication via a signaling gene pair^3^ (connexins or LR pairs), returning a list of cell type-signaling gene quadruplets with z-scores quantifying significance (**Fig. 1b,c**, Methods). A latent variable model (LVM) summarizes the resulting quadruplet set into interpretable modules and preference maps (Methods, **Fig. 1c**).

At the core is a statistical test for whether cell types *CT_i_* and *CT*_j_ communicate via genes *SG_k_* and *SG_l_*. CellWHISPER binarizes *SG* expression using a quantile-based threshold. Spatial information is represented by a *k* -nearest neighbors (KNN) graph^29^, from which a subgraph (“whisper network”) is derived to include edges between neighboring cells of types *CT_i_* and *CT*_j_ expressing genes *SG_k_* and *SG_l_* respectively (Fig. 1b). The whisper network edge count (*N_i_*_j*kl*_) quantifies the evidence for this quadruplet in the tissue sample. To assess significance, CellWHISPER shuffles locations among cells of same type (preserving cell-type expression and spatial distributions), recomputes *N_i_*_j*kl*_ and uses the resulting null to compute a z-score. For scalability, CellWHISPER derives closed-form expressions for the expectation and variance of the whisper network edge count under this null, avoiding explicit shuffling (Methods). The analytical z-scores agree closely with permutation-based z-scores (**Fig. 1f**) and are computable at orders-of-magnitude lower cost (**Fig. 1e**). The variance can be computed treating whisper edges as independent or accounting for edges that share a cell; the two give nearly identical rankings of quadruplets (Spearman ρ = 0.999; **Supp. Fig. 11**, Methods). In practice, this enables direct CCC inference on datasets comprising tens of thousands of cells and thousands of signaling gene pairs on standard hardware, without exceeding memory limits or incurring prohibitive runtimes. To assess statistical control for direct CCC, we applied the within-type randomization control described above. On downsampled Stereo-seq data (∼5,000 cells, 28 cell types, 367 LR pairs) competing tools (CellChat v2^22^, COMMOT^24^, SpaTalk^23^) reported similar numbers of significant quadruplets on real and randomized data whereas CellWHISPER reported far fewer quadruplets on randomized data, yielding an empirically estimated FPR <5%, underscoring its high specificity (**Fig. 1d**, Methods).

We focused initial validation on one of the best studied mechanisms of direct CCC: gap junctions (GJ). Gap junction communication provides a stringent stress test for direct CCC inference because they are purely contact-mediated and therefore particularly sensitive to spatial and expression confounders. GJ form when connexin protein in adjacent cells dimerize to establish intercellular channels^30^ (Fig. 1a). To evaluate the plausibility of using mRNA levels as proxies for connexin-mediated communication, we performed targeted spatial profiling in an induced pluripotent stem cell model^12^ and observed concordance between *Gja1* mRNA and Cx43 protein abundance, as well as substantial overlap between mRNA- and protein-derived whisper networks (**Supp. Figure 1, Methods**). In a Stereo-seq mouse brain dataset^27^ (Methods), aggregated whisper networks for connexin pairs were significantly larger than those from random genes (**Fig. 1g**, Mann-Whitney U test p-value 4.6e-8, Methods): each of 10 connexins yielded a whisper network of ≥8878 edges, while only 22/5000 (0.4%) random genes met this criterion (**Supp. Figure 2**).

### CellWHISPER identifies known and novel mechanisms of gap junction communication (GJC) in the mouse brain

To demonstrate the utility of CellWHISPER, we first applied it to discover GJC in mouse brain. From a computational perspective, GJC provides a stringent benchmark for direct CCC inference because it is purely contact-mediated and therefore maximally sensitive to spatial and expression confounders. GJC is believed to be widespread in the brain^31^ and has been implicated in neurodevelopment and diseases ^32–34^. Certain combinations of cell types and connexins are preferentially involved in brain GJC ^30,35^ and we sought to explore such a “connexin code”^16,28^. We analyzed a STEREO-seq mouse hemibrain dataset^27^, comprising of ∼50,000 cells across 28 cell types (CTs), using ten highly variable connexins as signaling genes (SG). (All 100 pairs of these genes were included in the SG compendium). CellWHISPER identified 1726 significant CT-SG quadruplets (z-score > 3, whisper network size > 30, **Fig. 2a**, **Supp. Data 1**) at an empirical FPR ∼1% estimated from randomized controls (**Supp. Figure 3**). These findings are stable across imputation parameters, binarization thresholds and neighborhood size (k) settings (Supp. Fig. 8) and under simulated errors in cell-type annotation (Supp. Fig. 9). This demonstrates that well-controlled inference over large quadruplet spaces is feasible at tissue scale (Fig 1e). Predictions were dominated by inter-glial GJC mechanisms (**Fig. 2b**).

**Figure 2.**
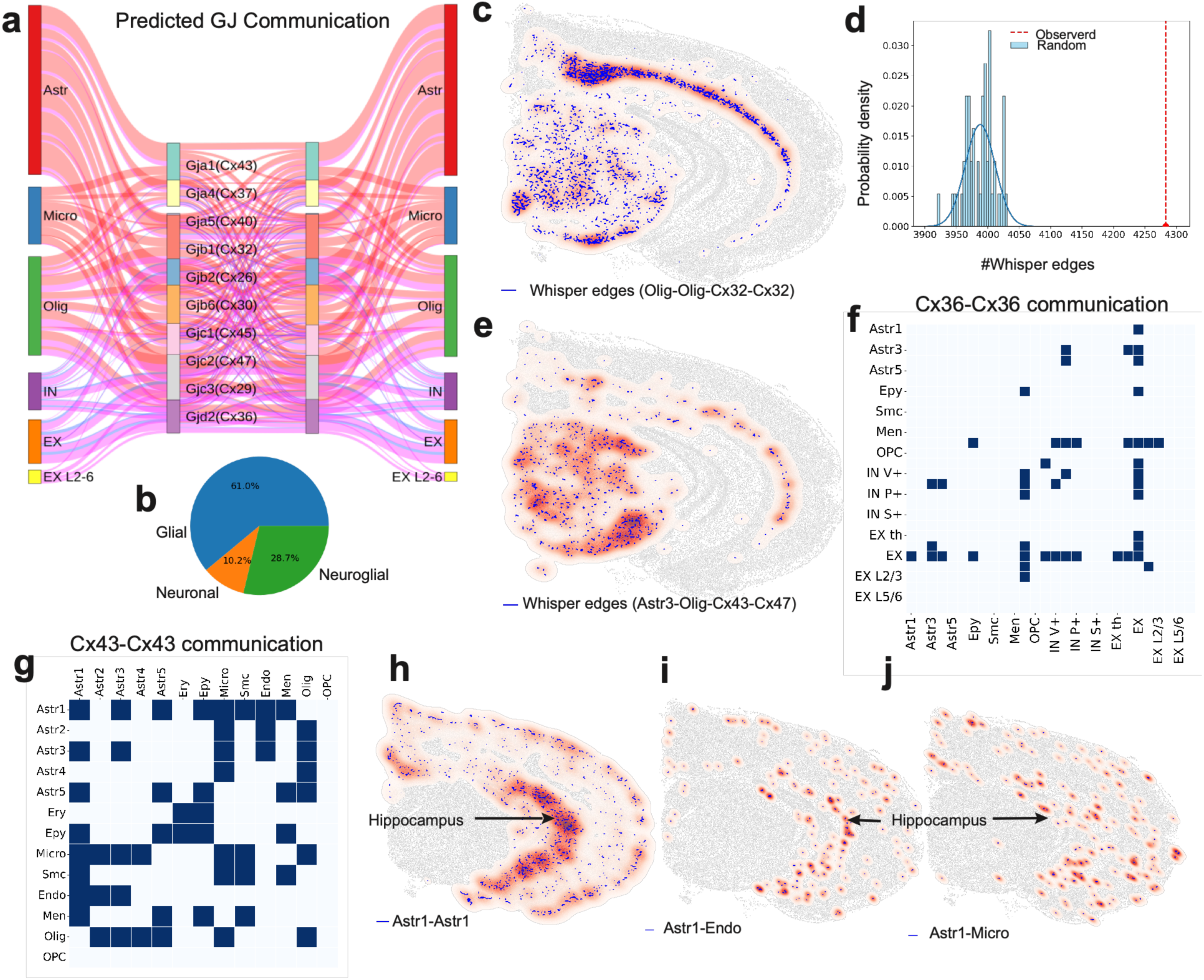
CellWHISPER Identifies Direct Gap Junction Communication in the Mouse Brain. **(a) Gap junction communication identified in mouse brain:** Sankey plot of the top 150 significant CT-SG quadruplets (a cell type pair communicating via a gap junction gene pair) ranked by whisper network size identified in Stereo-seq data (∼50K cells). For visual ease, “Astr” represents all astrocyte subtypes (Astr1-5), “IN” represents all interneuron subtypes and “EX L2-6” represents all excitatory neuron layers. (“Olig” = oligodendrocyte, “Micro” = microglia.) Lines indicate CellWHISPER-predicted communication involving specific cell type pairs (or aggregated groups such as Astr) and specific gap junction genes: red for glial-glial communication, blue for neuronal-neuronal communication, and pink for neuroglial communication. Line thickness reflects the number of significant quadruplets for each CT–SG pair, while node height reflects the total number of significant quadruplets associated with that cell type or signaling gene. (**b**) Pie chart of the three major classes of gap junction communication in the top 150 predicted CT-SG quadruplets shown in (a). (**c**) **Oligodendrocyte self-communication.** Whisper network of oligodendrocytes communicating via homotypic Cx32 gap junctions. Blue edges represent whisper network connections; red shading denotes kernel density smoothing at edge centers for visual clarity. (**d**) **Statistical significance of whisper edges.** Histogram of edge counts from randomized whisper networks (cell locations permuted within cell type), with fitted normal distribution (blue line). The red dashed line marks the observed number of edges in the whisper network shown in (c) which is significantly higher than expected under the null. Y-axis shows the probability density of randomized edge counts. (**e**) **Heterotypic astroglial communication.** Whisper network of astrocytes predicted to communicate with oligodendrocytes via heterotypic gap junction formed by Cx43 (astrocytic side) and Cx47 (oligodendrocytic side). (**f**) Cell types predicted to communicate using homotypic Cx36 gap junctions. Blue entries in heatmap denote significant cell type pairs (z score>3, whisper edge count > 30) found by CellWHISPER. Neuronal communication is predicted more frequently compared to glial communication. (**g**) Glial cell types predicted to communicate using Cx43-Cx43 gap junctions. Only glial communication is shown since it is predicted more frequently compared to neuronal communication (Supp. Fig. 6) (**h–j**) Astrocytic communication enriched in hippocampus. Whisper networks showing (h) astrocyte–astrocyte, (i) astrocyte–endothelial, and (j) astrocyte–microglia interactions mediated by homotypic Cx43 (Gja1) gap junctions

We examined the highest-confidence mechanisms from two complementary perspectives. Ranking quadruplets by whisper network size revealed that the top six represent inter-oligodendrocyte GJC via combinations of Cx47, Cx32, and Cx29 (**Supp. Data 1, Supp. Data 2**) consistent with established roles of Cx47 and Cx32 in oligodendrocytic coupling^36,37^. **Fig. 2c** illustrates homotypic Cx32 whisper network between oligodendrocytes, comprising 4,283 edges, far exceeding chance expectation (**Fig. 2d**, z-score = 13.09). The top predictions also included Cx29-mediated oligodendrocytic GJC. Although Cx29 transcripts are enriched in oligodendrocytes^38^, prior studies indicate that it primarily localizes to the adaxonal membrane rather than forming functional gap junctions^39^, highlighting a known limitation of transcript-based inference when subcellular protein localization is critical. The next highest-ranking quadruplets involve astrocyte-astrocyte GJC via Cx43, Cx30, and Cx26, connexins known to couple astrocytes into an astrocytic syncytium essential for homeostasis and adult neurogenesis^17,40–43^. We also identified high-confidence astrocyte-oligodendrocyte (A/O) communication via Cx43 on the astrocyte side and Cx47, Cx32 or Cx29 on the oligodendrocytic side. **Fig. 2e** illustrates *Astr3-*oligodendrocyte communication via Cx43-Cx47, consistent with prior reports supporting functional A/O GJ coupling^44^, though there is some unresolved debate about their functionality^43^.

Examining CellWHISPER results ranked by z-score rather than network size (**Supp. Data 3**) highlights additional mechanisms supported by fewer communicating cell pairs, which may thus be less well-studied in prior work. Among these, we identified GJC between excitatory neurons and other cell types, including thalamic excitatory neurons, oligodendrocytes, and astrocytes (**Supp. Data 3**). Maubecin et al.^45^ provided direct evidence of functional neuro-glial coupling using heterotypic gap junctions in rodents. Gap junctions between excitatory neurons are known to be primarily mediated by Cx36 (Gjd2)^46,47^ and extensively studied in rodent models^48–51^. As shown in **Fig. 2f**, excitatory neurons are a major site of homotypic Cx36-mediated communication predicted by CellWHISPER.

### CellWHISPER identifies homotypic Cx43 gap junctions as a mechanism underlying astrocytic syncytium

Cx43 is the most studied connexin, with key roles in differentiation and homeostatic regulation^52–54^. We examined the complete map of cell type pairs predicted by CellWHISPER to participate in Cx43-mediated GJC (**Fig. 2g**), finding a predominance of glial over neuronal communication. This analysis provides a focused validation setting, as Cx43-mediated interactions are well characterized. Astrocyte subpopulation Astr1 is predicted to communicate with other astrocytes, as well as endothelial, microglia, oligodendrocytes and ependymal cells. This aligns with known biology: Cx43 mediates astrocyte-oligodendrocyte-ependymal cell interactions forming an isopotential syncytium ^53,55–57^. Predicted astroglial interactions exhibited a higher network density in the hippocampus (**Figs. 2h-j**). Inter-astrocyte GJC (z-score 18.13, **Fig. 2h**) in the hippocampus has been documented to be Cx43-mediated. Astrocyte-endothelial GJC (z-score 10.94, **Fig. 2i**) has been implicated in maintaining blood–brain barrier integrity^58–60^. Additionally, CellWHISPER predicted Cx43-mediated communication between microglia and astrocytes (**Fig. 2j**) as well other microglia. Indeed, recent studies suggest that microglia upregulate Cx43 under pathological or inflammatory conditions and form functional gap junctions – with each other and with astrocytes^61–63^.

To test these computationally inferred predictions, we performed immunofluorescent staining on mouse coronal brain sections for Cx43 together with markers of astrocytes (GFAP), endothelial cells (CD31), and microglia (IBA1) (**Fig. 3a, b**; Methods). All channels showed robust signal across brain regions (**Fig. 3b**). Distinct Cx43 co-localization was evident at astrocyte–astrocyte, astrocyte–endothelial, astrocyte–microglial, and microglial–microglial interfaces (**Fig 3b,c**). Quantitative analysis showed higher co-localization of Cx43 with cell-type markers in the hippocampus compared to cortex (**Fig. 3d**), with astrocytic contacts enriched >5-fold. Representative examples of colocalization of astrocytes, microglia and Cx43 in the hippocampus are shown in **Fig. 3c**. Notably, these experimentally supported Cx43 interfaces (Astrocyte– Astrocyte, Astrocyte–Endothelium, and Astrocyte–Microglia) are prioritized among the highest-ranked CellWHISPER predictions, enabling targeted experimental follow-up. By contrast, CellChat v2 and COMMOT, run on the same dataset and connexin pairs, assign their minimum p-value to 12,358 and 9,680 quadruplets respectively, including these interfaces, limiting their usefulness for prioritizing specific mechanisms. To recover five reference interactions (**Supp. Fig. 10**), a user would need to screen 620 CellWHISPER predictions, compared with 12,358 for CellChat v2 and 9,680 for COMMOT. Together, these observations support the ability of CellWHISPER to recover known, region-specific contact-mediated communication patterns using a rigorous confounder-aware framework.

**Figure 3.**
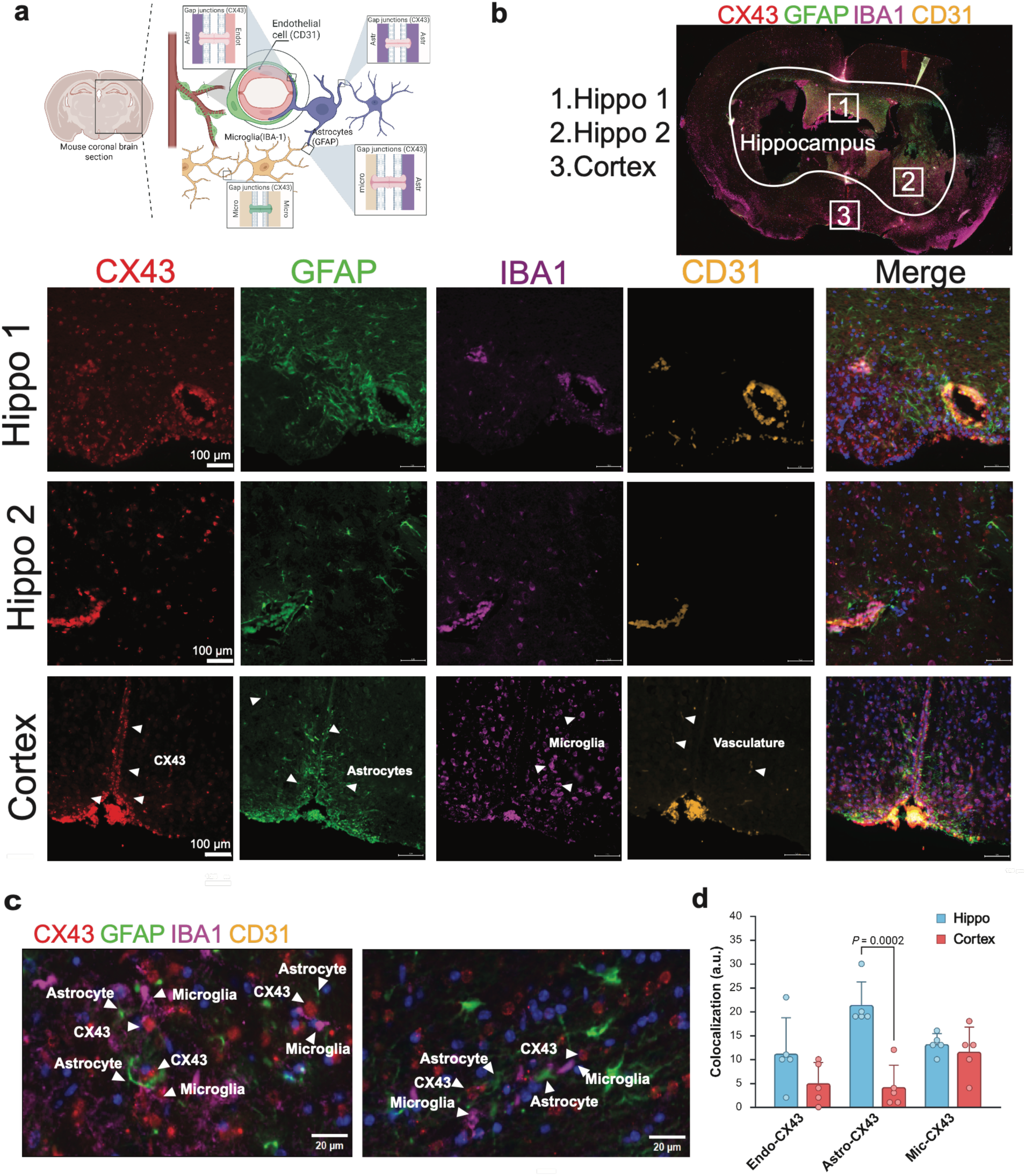
Spatially resolved analysis of Cx43-mediated cellular crosstalk across brain regions. (**a**) Schematic of gap junction–mediated interactions between astrocytes (GFAP), microglia (IBA1), and endothelial cells (CD31) via Cx43 (created with BioRender.com). (**b**) Immunofluorescence of coronal mouse brain section stained for Cx43, GFAP, IBA1, and CD31. Hippocampal boundaries were defined using the Allen Mouse Brain Atlas. Regions of interest (ROI) within hippocampus (Hippo1, Hippo2) and cortex were outlined for higher-resolution imaging. Scale bar, 100 µm. (**c**) Single-cell–resolution images of hippocampal ROIs showing colocalization of astrocytes and microglia with Cx43. (**d**) Quantification of Cx43 colocalization across hippocampus and cortex. We selected 10 regions (478 × 262 pixels each) within and outside the hippocampus and quantified Cx43 colocalization with astrocytes, endothelial cells, and microglia. Colocalization was defined as instances where Cx43 puncta were positioned between or overlapping the indicated cell types, yielding 30 values (one per cell type per region) and a total of 333 interactions. Two-way ANOVA revealed a significant enrichment of astrocyte–Cx43 interactions in the hippocampus compared to the cortex (p = 0.0002).

### CellWHISPER identifies direct ligand-receptor (LR) communication

Direct LR communication plays a fundamental role in organizing local tissue architecture^21^. We next used LR interactions as a test of generality of the CellWHISPER analytic framework. Unlike gap junctions, LR signaling spans diverse functional classes (adhesion, guidance, immune and ECM), while still requiring spatial proximity for direct interactions. To identify such interactions in the mouse brain, we re-analyzed STEREO-seq data using 367 curated cell-cell contact LR pairs from the CellChat database^5^ (Methods). CellWHISPER identified 4743 significant CT–SG quadruplets (z-score > 4, whisper network size > 30) (**Fig. 4a, Supp. Data 4**), with an empirically estimated FPR < 1%, spanning inter-glial, neuron-glia and neuron-neuron communication (**Fig. 4b**).

**Figure 4.**
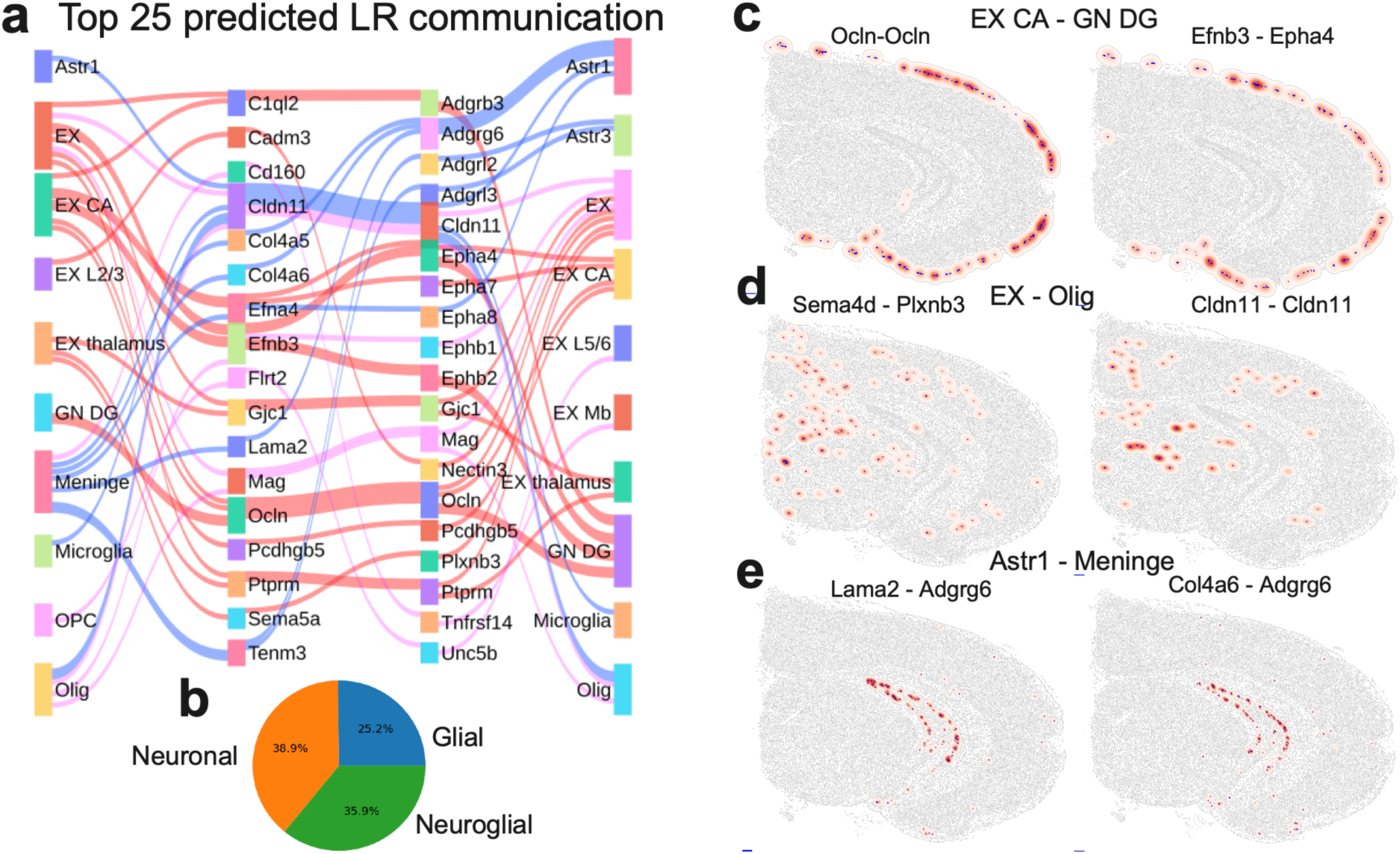
CellWHISPER Identifies direct Ligand-Receptor communication in the mouse brain. **(a)** Ligand–receptor communication mechanisms. Sankey plot showing the top 25 significant CT–SG quadruplets **ranked by z-score** (z-score > 4, whisper network size > 30) identified using 367 curated ligand–receptor pairs. Randomized control analysis estimated a false positive rate (FPR) < 1% for this compendium. **(b)** Communication classes. Distribution of major interaction types, including neuron–neuron, neuron–glia, and inter-glial communication. **(c–e)** Six representative whisper networks (two per panel, left/right). Selected ligand–receptor–mediated cell–cell interactions illustrating key modes of direct communication: (c) excitatory CA3 neurons (EX CA) and dentate gyrus granule neurons (GN DG) communicating via Efnb3-Epha4 (right) and tight junction Ocln (left); (d) excitatory neuron-oligodendrocyte interactions involving *Cldn* (right) and semaplexin family member (left); and (e) meningeal-astrocyte communication via extracellular matrix components (*Lama2*, *Col4a1–6*, *Adgrg6*).

The highest confidence predictions revealed communication between CA3 excitatory neurons (EX CA) and dentate gyrus granule neurons (GN DG) mediated by LR pairs of the mossy fiber pathway^64^ (**Supp. Data 5**). These include multiple ephrin/Eph family members, C1ql2/3-Adgrb3 and homotypic interactions of occludin, protocadherin and claudin (**Supp. Data 5, Fig. 4c**). Ephrin–Eph and C1ql2/3–Adgrb3 (BAI3) interactions, both implicated in mossy fiber–CA3 synapse organization^65–67^, ranked among the highest-confidence predictions, alongside multiple adhesion-related pairs (including claudins/occludin). These findings are consistent with known hippocampal wiring programs^68^ and support the biological plausibility of CellWHISPER’s proximity-constrained LR inference.

CellWHISPER also predicted LR-mediated communication between oligodendrocytes and excitatory neurons, involving homotypic MAG, claudin and semaplexin interactions (**Supp. Data 6, Fig. 4d**), consistent with known roles for these mechanisms in axon–glia recognition, myelin stability^69^, and activity-dependent modulation of myelination^70^. Additionally, we observed interactions between astrocytes and meningeal cells, notably involving extracellular matrix components (ECM) such as Lama2, type IV collagens (Col4a1–6), Adgrg6 (**Supp. Data 7, Fig. 4e**). These examples illustrate that CellWHISPER supports discovery across multiple LR functional categories while maintaining stringent specificity for direct CCC.

### Latent Variable Modeling (LVM) infers general patterns of communication mechanisms

We used CellWHISPER’s “LVM” to infer higher-order patterns among significant quadruplets (Methods). Unlike approaches that first collapse quadruplet-level information into CT x SG or CT x CT matrices^5,6^, an aggregation step that can be information-losing; CellWHISPER directly models quadruplets to characterize communication profiles of cell types and signaling genes. Specifically, the LVM models the presence or absence of statistically supported CT–SG quadruplets using a shared low-dimensional embedding space for cell types and signaling genes (Methods). Under this model, a quadruplet is more likely when each cell type exhibits higher affinity for its corresponding signaling gene, yielding embedding geometry that captures CCC preferences. The learned embeddings yield (1) CT–SG preferences, (2) CT–CT similarity, and (3) SG–SG similarity. Model parameters are estimated via constrained log-likelihood maximization. **Fig. 5b** shows that the optimization can accurately infer cell type similarity on simulated data (Methods).

**Figure 5.**
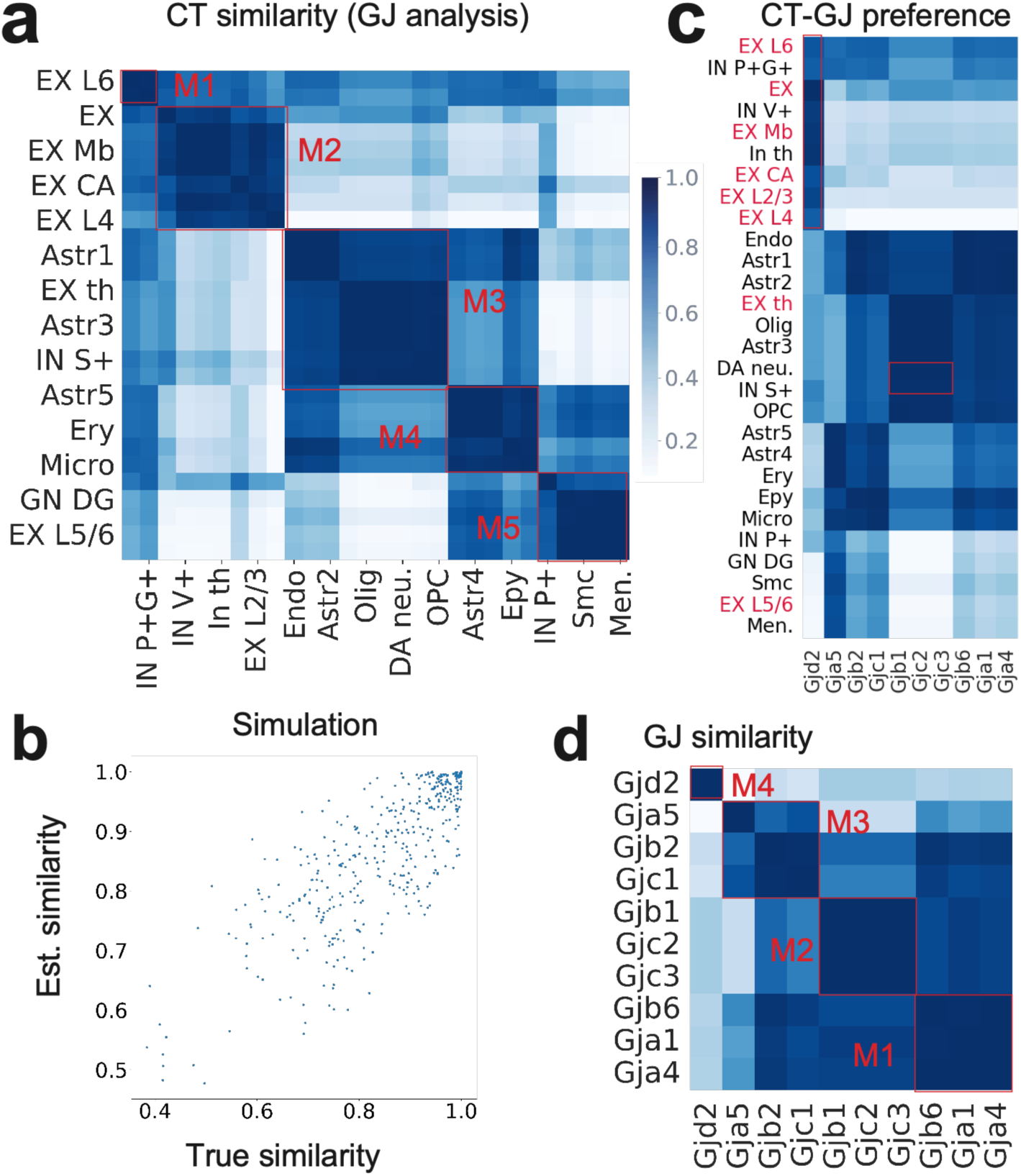
LVM infers module of signaling cell types and signaling genes. (**a**) Cell type similarity matrix. Cosine similarity of latent cell-type embeddings derived from the LVM routine of CellWHISPER, applied to gap junction communication in the Stereo-seq mouse brain dataset. Five dominant modules of cell types emerge, reflecting recurrent patterns of gap junction communication across the tissue. (**b**) Recovery of cell-type similarity on simulated data. Constraint optimization in latent variable model identifies underlying cell type pair similarity, resolving higher-order structure in the data. Each point is a cell-type pair; x-axis, true cosine similarity of the embeddings used to simulate the data; y-axis, cosine similarity of the embeddings estimated by the LVM (Methods). (**c**) Cell type–gap junction preferences. Heatmap showing associations between cell type modules and gap junction gene modules. Distinct modules exhibit preferential coupling, e.g., Module M1 and M2 are enriched in excitatory neurons and prefer Gjd2 (GJ module M4). DA neurons and IN S+ prefer GJ module M2. **(d)** Cosine similarity of latent embeddings across gap junction genes, revealing four dominant connexin modules.

Applying LVM to GJ results in mouse brain Stereo-seq data revealed five dominant cell type modules in CT-CT similarity map (**Fig. 5a, Supp. Data 8**) whereas a baseline method performing naïve aggregation of quadruplet tensor did not reveal modular structure (**Supp. Figure 4**). LVM also identified four GJ modules in SG-SG similarity map (**Fig. 5d**, Supp. Data 8). Cell type modules showed preferences for specific GJ modules in the CT-SG preference matrix (**Fig. 5c**). For example, excitatory neuron subtypes in cell type modules M1/M2 (EX Mb, EX L2/3, EX CA) prefer GJ module M4 consisting of Gjd2 (Fig. 5c,d). Notably, DA neurons (excitatory or inhibitory depending on context^71^), clustered with inhibitory Sst+ interneurons^72^ in cell type module M3 (Fig. 5a, Supp. Data 8) and shared preference for GJ module M2 (Fig. 5c), consistent with dye-coupling evidence of DA–interneuron electrical synapses^73^ and suggesting candidate GJ mediating these synapses.

Next, we applied LVM to LR interactions (Fig. 4), identifying four dominant cell-type modules (**Supp. Fig. 12a, Supp. Data 9)**. Excitatory neurons in cortical layers and CA region co-cluster, whereas thalamic excitatory neurons lie in different module (**Supp. Fig. 12a,b**). This aligns with known features of excitatory neurons in the neocortex and hippocampus (of which CA is part), including laminar structure^74,75^ and shared neuromodulatory response^76^, different from excitatory neurons in thalamic neurons^77^. Analysis of ligand similarity map (**Supp. Fig. 12e**) identified five dominant modules, of which one (M1) is enriched for cell adhesion, ECM and immune modulators (**Supp. Data 9**). This ligand module preferentially associates with cell type module M1 (**Supp. Fig. 12c),** enriched for glial cell types located near the brain periphery, such as ependymal cells, meninges, and smooth muscle cells (Supp. Fig. 12a), suggesting a specialized glial–meningeal signaling axis potentially involved in barrier maintenance and neuroimmune communication. Another ligand module (M5, Supp. Data 9) is enriched for axon guidance and synaptic signaling genes^78,79^, and prefers cell type module M2 (**Supp. Fig. 12c**), enriched in excitatory neurons in the cortical layers and hippocampus, consistent with their known role in neuronal cell-cell communication^80^. Similarly, we found five dominant modules in receptor similarity map (**Supp. Fig. 12f**, Supp Data 9) having preference for specific cell type modules (**Supp. Fig. 12d**). Together, LVM captures dominant communication patterns across GJC and LR signaling.

### Contrasting CellWHISPER results on an Alzheimer’s disease model with wild-type brain

Gap junction-mediated communication (GJC) supports central nervous system homeostasis and is implicated in neurodegenerative diseases, including Alzheimer’s disease (AD)^30,81,82^. Under normal conditions, GJC supports metabolic and synaptic function, especially through astrocytic networks^62,83^ whereas in AD, aberrant gap junctions and hemichannel activity contributes to neuroinflammation and neurodegeneration^30,81,82^ ^30^. From a computational perspective, this analysis tests whether CellWHISPER can identify condition-associated rewiring using the same statistically well-controlled, proximity-constrained quadruplet framework, while limiting false positives arising from differences in cell-type composition, spatial organization, or expression. To systematically interrogate these dynamics, we used CellWHISPER to decode GJC networks altered in AD.

We ran CellWHISPER separately on Xenium mouse brain data from wild-type and TgCRND8 mice^84^ (one section per condition, aged >13 months, Methods) using three annotated GJ genes. After cross-sample integration to harmonize cell-type labels (Methods), we compared quadruplets passing thresholds in both conditions (shared) or in one condition only (Methods). This revealed 60 quadruplets (z-score > 4, whisper network size > 30, **Fig. 6a, Supp. Data 10**) in at least one condition. The twenty shared quadruplets were enriched for astrocytic (13/20) and oligodendrocytic communication (9/20) (**Fig. 6b**), including Astr1-Olig1 homotypic Cx43 (**Fig. 6c**). Despite known Cx43 upregulation near Aβ plaques, prior work suggests structural preservation of astrocytic GJC in AD brain^85^; consistent with this, CellWHISPER identifies conserved GJC mechanisms despite change in signaling gene expression.

**Figure 6.**
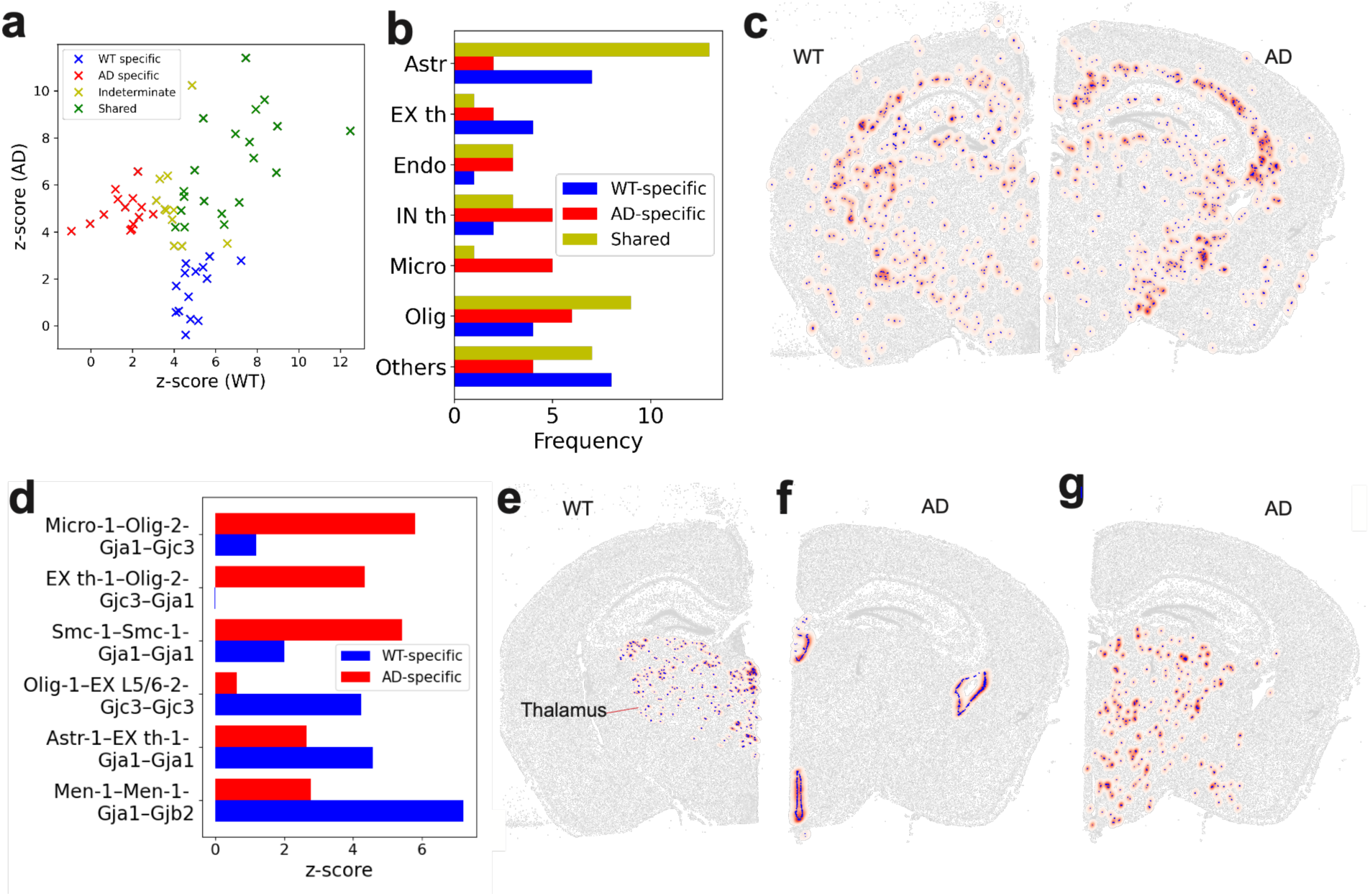
Comparison of gap junction communication between wild-type and Alzheimer’s disease (AD) model. (**a**) Scatter plot of 60 high-confidence CT–SG quadruplets (z-score > 4, whisper network size > 30 in at least one condition) showing their z-scores in wild-type (WT) and TgCRND8 (AD) Xenium datasets for Gja1, Gjb2, and Gjc3. Quadruplets are classified as shared, condition-specific (detected in one condition only), or indeterminate. (**b**) Bar plot showing the frequency of each cell type across the 60 quadruplets. Astrocytes and oligodendrocytes are most enriched in shared interactions, microglia dominate AD-specific interactions, and astrocytes are more frequent in WT-specific interactions. (**c**) Whisper networks of Astr1–Olig1 communication via homotypic Cx43 in WT (left) and AD (right), showing preserved symmetry between conditions. (**d**) Selected condition-specific quadruplets, showing z-scores in WT versus AD. Top three WT-specific and AD-specific interactions were chosen based on whisper network size. (**e**) Whisper network of Astr1–Ex th1 communication via homotypic Cx43 in WT. (**f**) Whisper network of Smc1– Smc1 (smooth muscle cell self-communication) via homotypic Cx43 in AD, highlighting potential vascular remodeling. (**g**) Aggregated microglial whisper network in AD, constructed by taking the union of all AD-specific quadruplets involving microglia, revealing enrichment in thalamus and hypothalamus regions.

We further observed 15 TgCRND8-only and 14 wild-type-only quadruplets (Fig. 6a,d, **Supp. Data 10**), including reduced astrocyte-thalamic excitatory neuron GJC via homotypic Cx43 (Gja1) (**Fig. 6e**). The corresponding whisper network (Fig. 6e) shows enrichment of this communication in the WT thalamus. In contrast, TgCRND8-only mechanisms included increased smooth muscle cells homotypic Gja1 (Fig. 6d,f), potentially reflecting vascular remodeling^86,87^. TgCRND8-only quadruplets were enriched for microglial interactions (5/15 vs. 0/14 in WT), and WT-only quadruplets for astrocytes (6/14 vs. 2/15 in TgCRND8) (**Fig. 6b**). These trends align with existing literature: astrocyte networks maintain homeostasis in healthy brain^88^, while microglial GJC becomes more prominent during AD neuroinflammation^30,89,90^. Aggregated microglial whisper network (**Fig. 6g**) showed enrichment in thalamus and hypothalamus. Connexin alterations have been reported in thalamic regions in AD^81^ while microglial changes in thalamus are linked to cognitive impairment after cortical brain injury^91^.

### Application of CellWHISPER to tissues with spatial gradients

Many tissues are organized along spatial gradients, such as the crypt-to-surface axis of the colonic epithelium or the stratified layers of the epidermis, along which cell states and gene expression change continuously. Such gradients pose a particular challenge for CCC inference, since cells of the same type and their signaling genes are co-localized by construction, and zonal co-expression can be mistaken for communication. Because CellWHISPER randomizes cell identities within each cell type, spatial organization encoded by the cell-type labels, including zones and layers, is preserved under the null distribution, and a quadruplet is called significant only when co-expression at cell contacts exceeds what this organization predicts. We tested this on two human tissues profiled with Xenium, non-diseased colon (425-gene panel) and skin (5K panel), annotating spatially defined cell states as part of the cell-type labels (crypt-base to surface epithelial types in colon; basal and three suprabasal keratinocyte layers in skin). We tested every cell-cell contact pair from CellChatDB present on each panel (Methods, **Supp. Data 11 and 12**).

In colon, a per-cell zonation score recovered the expected crypt-to-surface gradient across a 58,125-cell region (Fig. 7a). Among the 1,491 quadruplets tested (9 contact pairs on the panel), the top-ranked quadruplet was *CEACAM1-CEACAM5* between enterocytes (z-score 42.5). Its whisper network traces the surface epithelium lining each crypt (Fig. 7b), consistent with the surface-restricted expression of *CEACAM*s in colonic epithelium. In the same region, *FCER2* (CD23)-*CR2* (CD21) between B cells was significant (z-score 12.1) with a whisper network confined to lymphoid follicles of the lamina propria (Fig. 7c), a compartment that lies off the crypt axis altogether. Zonal co-expression alone did not lead to significant CCC calls: *CEACAM1-CEACAM5* contacts among BEST4+ epithelial cells, which express both genes almost uniformly within the same surface zone, were not significant (z-score 0.27), because the expected number of co-expressing contacts under the null distribution already reflects the co-localization of these cells.

**Figure 7.**
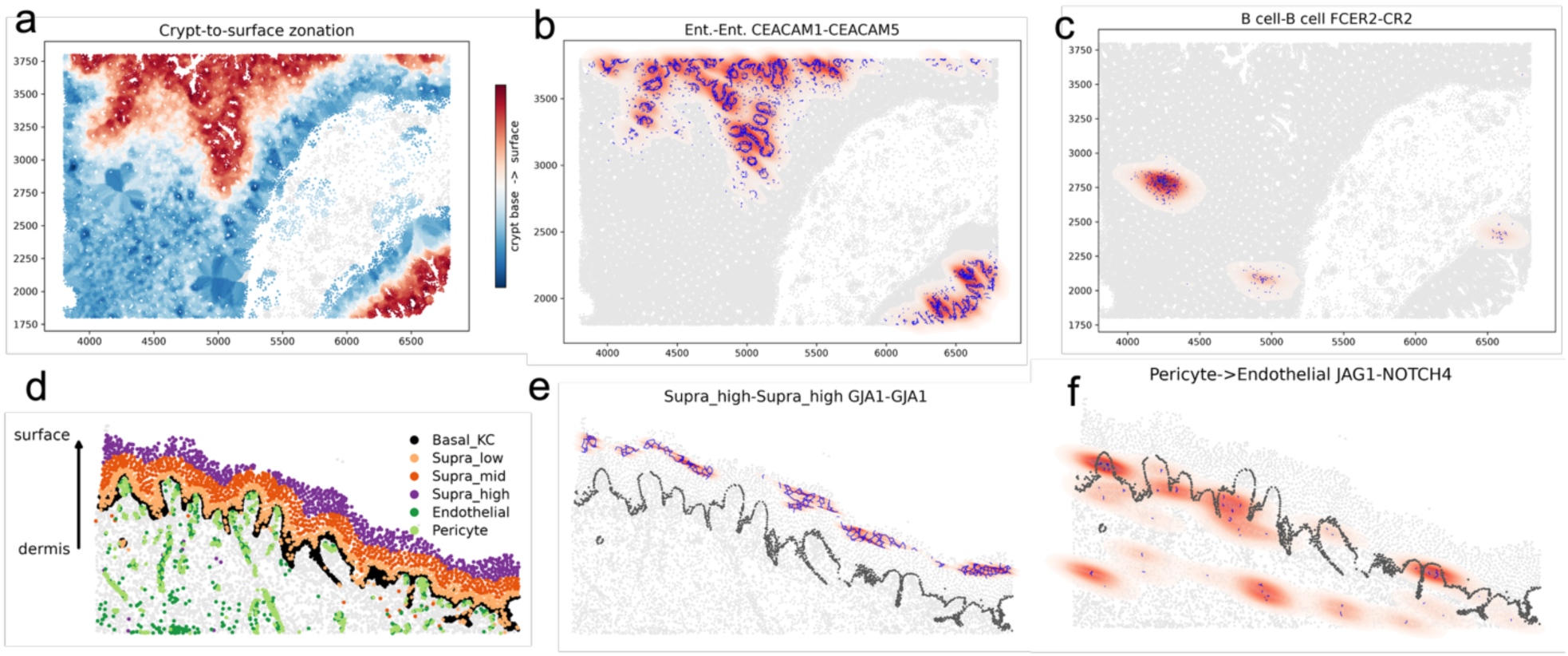
CellWHISPER identifies contact-mediated communication in spatially zonated human tissues. (**a**) Crypt-to-surface zonation in a 58,125-cell region of human colon (Xenium). Each cell is colored by a positional zonation score, the mean surface-marker minus crypt-base-marker expression of its ten nearest epithelial cells (blue, crypt base; red, surface); cells far from the epithelium are shown in grey. (**b**) Whisper network for Enterocyte–Enterocyte communication via *CEACAM1-CEACAM5*, the top-ranked quadruplet among all CellChatDB contact pairs on the panel (z-score 42.5, 5,684 whisper edges). Blue edges are whisper edges; red shading is the kernel density of edge midpoints, as in Fig. 2c. The network traces the surface epithelium lining each crypt. (**c**) Whisper network for B cell–B cell communication via FCER2-CR2 (z-score 12.1, 433 edges) in the same region, confined to lymphoid follicles of the lamina propria. (**d**) Cell types in the epidermis and underlying dermis of a human skin section (Xenium Prime 5K; 12,114 cells). The arrow indicates the differentiation gradient from the dermal-epidermal junction to the surface. (**e**) Whisper network for homotypic Cx43 (GJA1) gap-junction communication between keratinocytes of the outer suprabasal layer (z-score 13.0, 464 edges), forming a band along the surface of the epidermis. (**f**) Whisper network for JAG1-NOTCH4 communication from pericytes to endothelial cells (z-score 14.6, 111 edges), the top-ranked quadruplet of the skin scan, forming discrete foci at varying distances from the epidermis.

Analysis of skin tissue faced a sharper gradient, from the basal layer at the dermal-epidermal junction to the outer epidermis (Fig. 7d). Of 10,773 quadruplets tested (145 contact pairs on the panel), 65 quadruplets were found significant (**Supp. Data 12**). Along the gradient, homotypic Cx43 (GJA1) gap junctions between keratinocytes of the outer suprabasal layer were significant (z-score 13.0, Fig. 7e), where Cx43 coupling of differentiating keratinocytes is well established, whereas the same gene between basal keratinocytes, in which it is expressed uniformly, was not (z-score 1.1). Off the gradient, the top-ranked quadruplet of the entire scan was *JAG1-NOTCH4* from pericytes to endothelial cells (z-score 14.6), a canonical circuit of mural-endothelial signaling, whose whisper network forms discrete foci at varying distances from the epidermis rather than a band along it (Fig. 7f). *DLL4-NOTCH3* and *JAG1-NOTCH3* from endothelial cells to pericytes were likewise among the top predictions (Supp. Data 12). Together, the colon and skin analyses show that CellWHISPER can recover contact-mediated communication both along and independent of tissue gradients, without being confounded by zonal co-expression.

## DISCUSSION

Direct cell–cell communication shapes tissue homeostasis and disease, yet existing spatial transcriptomics tools conflate genuine signaling with cell type co-localization, limiting their utility for rigorous inference. CellWHISPER addresses this by explicitly accounting for confounding effects of gene-expression and spatial organization, CellWHISPER markedly reduces FPR relative to competing tools. Two additional computational features broaden practicality and interpretability: an analytical null that avoids explicit permutation testing and enables scalable inference across large tissues and signaling compendia, and a quadruplet-native latent variable model that extracts higher-order structure without collapsing interactions into information-losing pairwise summaries. The framework is agnostic to the signaling compendium (connexins, curated LR pairs, or user-defined contact genes) and scales to tens of thousands of cells, enabling routine application to emerging high-resolution spatial platforms.

We demonstrated CellWHISPER on mouse brain spatial datasets as validation settings spanning two major classes of direct CCC: gap junctions and contact-dependent ligand–receptor interactions. CellWHISPER recovered canonical glial networks and generated a large-scale map of predicted connexin-mediated interactions (“connexin code”). Immunostaining confirmed Cx43 colocalization at astrocyte–astrocyte/endothelial/microglial interfaces. LR analysis uncovered adhesion/synaptic programs, including the mossy fiber pathway and a glial–meningeal signaling axis. Comparison of an AD mouse model identified a shift toward microglial-associated GJC and region-specific remodeling in the thalamus and vasculature, consistent with emerging models of neuroinflammatory progression. Beyond brain, application to human colon and skin showed that in tissues with strong spatial gradients CellWHISPER recovers contact circuits both along the gradient and independent of it, without being unduly confounded by zonal co-expression of ligand-receptor genes (Figure 7).

Several limitations motivate future work. CellWHISPER inference relies on mRNA as a proxy for membrane proteins, and post-transcriptional regulation or subcellular localization can yield false positives. CellWHISPER detects enrichment of co-expressing cell pairs at contact, which prioritizes a mechanism for experimental follow-up rather than establishing functional coupling. Since it operates on segmented cell bodies, communication through dendrites and axons far from the soma, a substantial fraction of direct signaling in brain, is beyond its reach, as for any method based on segmented spatial transcriptomics. Segmentation errors in ST^92^ may also propagate to CCC inference, a limitation shared across current methods. CellWHISPER prioritizes specificity through a confounder-aware null model, which may come at the cost of sensitivity. At the same time, it excels at detecting communication events mediated by a smaller number of cell pairs. The null distribution used in its testing preserves the spatial organization of given cell-type labels, so its statistical guarantees are conditional on the annotation: in tissues with gradients, spatially defined states such as crypt zones or epidermal layers should be part of the labels, as in Figure 7. The wild-type versus TgCRND8 comparison rests on one section per condition; establishing disease-associated changes in communication will require biological replicates. Significance calls are robust to modest levels of annotation error (Supp. Fig. 9) and to preprocessing and graph settings (Supp. Fig. 8). Extensions to lower-resolution platforms (e.g. Visium) may be possible via deconvolution approaches, and integrating pathway information could improve robustness of whisper networks. Other spatial tools address related questions at different levels of resolution: SpatialDM^93^ prioritizes ligand-receptor pairs by their spatial co-expression, while DeepTalk^94^ and CellNEST^95^ score communication between individual cells, the latter with a focus on relay networks. These outputs are complementary to CellWHISPER’s cell-type-pair resolution and do not provide a null-referenced significance call, so they were not included in our benchmark.

In summary, CellWHISPER establishes a robust paradigm for inferring direct signaling from ST data, combining stringent statistical control with interpretable latent structure, to enable mechanistic dissection of tissue organization and for identifying disease-specific communication in development, homeostasis, and disease.

## METHODS

### Overview of CellWHISPER

CellWHISPER infers direct cell–cell communication (CCC) from spatial transcriptomics (ST) data. The input consists of gene expression profiles, spatial coordinates, and cell-type annotations for individual cells, together with a precompiled compendium of signaling gene (SG) pairs, such as connexins or ligand–receptor (LR) pairs. SG compendia can be obtained from existing resources or provided by the user as a custom list.

For each candidate combination of a cell-type pair (*ct_i_*, *ct*_j_) and a signaling gene pair (*sg_k_*, *sg_l_*), CellWHISPER tests whether the two cell types exhibit evidence of direct communication mediated by the signaling genes (Fig. 1b). Statistically supported interactions are reported as CT–SG quadruplets, each associated with a z-score quantifying significance (Fig. 1c). CellWHISPER includes two analysis modules (Fig. 1c): (i) A statistical testing module identifies significant CT– SG quadruplets, and (ii) A latent variable model (LVM) summarizes the collection of significant quadruplets into interpretable preference and similarity maps, including CT–CT, CT–SG, and SG– SG relationships.

### Whisper network construction

Gene expression is binarized using a per-gene quantile threshold (75th percentile across all cells by default), yielding a binary indicator of whether a signaling gene is expressed in a given cell. A per-gene quantile places genes of different dynamic range and sparsity on the same scale, so that the per-cell-type expression probabilities used in the null distribution are comparable across genes; it also keeps the null tractable, since these probabilities are simple fractions of cells, and suppresses low-level expression noise. For a sparse gene the 75th percentile is zero, and a gene is then considered expressed in every cell in which it is detected.

Spatial proximity between cells is represented using a k-nearest neighbors (KNN) graph constructed from cell coordinates (k = 5 unless otherwise specified), symmetrized so that an edge is kept only if each cell is among the other’s k nearest neighbors; the graph is therefore undirected and has no self-edges. This graph defines the set of potential physical contacts used for CCC inference. For datasets with irregular tissue geometry, distance-based thresholds should be used instead of fixed neighbor counts.

For each CT–SG quadruplet (*ct_i_*, *ct*_j_, *sg_k_*, *sg_l_*), a “whisper network” is derived by retaining only edges connecting neighboring cells of types *ct_i_* and *ct*_j_ that express *sg_k_* and *sg_l_*, respectively (Fig. 1b). Quadruplets are directional, with *sg_k_* assigned to *ct_i_* and *sg_l_* to *ct*_j_, so (*ct_i_*, *ct*_j_, *sg_k_*, *sg_l_*) and (*ct*_j_, *ct_i_*, *sg_k_*, *sg_l_*) are tested separately; for *i* = *j* each pair of neighbouring cells is counted once. The number of edges in this subgraph, denoted *N_i_*_j*kl*_, quantifies spatially supported evidence for the corresponding communication mechanism.

### Statistical test for CT–SG quadruplets

To assess statistical significance, CellWHISPER constructs a null model by permuting cell identities within each cell type. This preserves cell-type–specific gene expression distributions and spatial organization while disrupting spatial proximity between signaling cells of different types. The whisper network is recomputed under the null, yielding a null distribution for *N_i_*_j*kl*_, from which a z-score is computed.

CellWHISPER derives the expectation and variance of *N_i_*_j*kl*_ under this randomization in closed form (below); permutations are not performed in practice and were used only to verify the analytical null (Fig. 1f). The expectation is exact. The variance can be computed in one of two modes: an independent-edge (iid) mode that treats whisper edges as independent Bernoulli variables, and a shared-cell (non-iid) mode that additionally accounts for pairs of edges sharing a cell. The two give nearly identical rankings of quadruplets, the shared-cell variance being slightly larger and therefore more conservative (Supp. Fig. 11). The iid mode has negligible cost beyond the expectation, whereas the shared-cell mode requires the degree of every cell in each cell-type-pair subgraph.

### Analytical null for the statistical test of a quadruplet

Let *Z_u_* be a Boolean random variable indicating the presence of a whisper edge on proximal cell pair *u* for the quadruplet *i*, *j*, *k*, *l*. Let *N_i_*_j_ be the number of proximal pairs between cell types *i* and *j*. We derive the expectation and variance of whisper edge count ∑*_u_ Z_u_* under within-cell-type randomization. The probability that edge *u* is a whisper edge (i.e. *Z_u_* = 1) is the product of the probabilities that gene *k* is expressed in a cell of type *i* and gene *l* is expressed in a cell of type *j*. Gene expression is determined by a global percentile threshold across all cells. Thus:

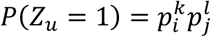

where 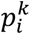 is the fraction of cells of type *i* in which gene *k* exceeds its binarization threshold and 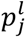 correspondingly for gene *l* in cell type *j*.

### Independent-edge mode (iid)

Treating the *Z_u_* as independent Bernoulli variables, the count follows a Binomial distribution with *N_i_*_j_ proximal pairs:

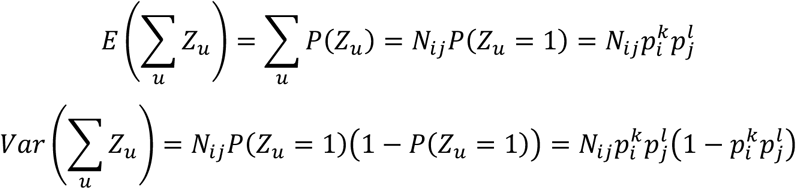

### Dependence-aware mode (non-iid)

Proximal pairs that share a cell are not independent. The expectation is unchanged by linearity, the variance is:

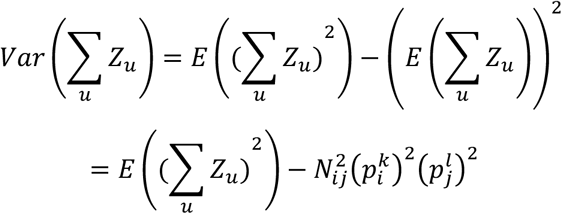

Expanding the first term,

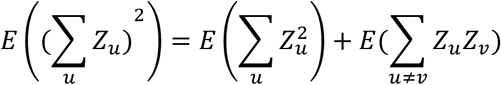

Since, 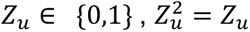, so the first sum reduces to E(∑ *Z*) = *N_ij_ p_i_ p_j_*. In the second term each component *E*(*Z_u_Z_v_*) is the product of expression probabilities over the distinct cells involved. For *i* ≠ *j*, pairs (*u*, *v*) fall into three cases:

- no shared cell: *Zu*, *Zv* are independent, 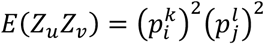
- shared cell of type *i* but distinct cells of type 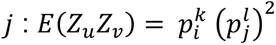
- shared cell of type *j* but distinct cells of type *i* : 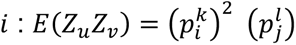

Counting: Let *C_i_* be the number of ordered pairs of proximal edges that share a cell of type *i*, and *C*_j_ likewise for type *j*. For a cell *c* of type *i* (denoted by “*t_c_* = *i*”) with degree *d_c_* (its number of proximal neighbors of type j), there are *d_c_*(*d_c_* − 1) ordered edge pairs sharing *c*, so *C_i_* = ∑*_c_*_:*t*=*i*_ *d_c_*(*d_c_* − 1) and likewise *C*_j_ = ∑*_d_*_:*td*=j_ *d_d_*(*d_d_* − 1) for cells of type *j*. With 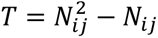 ordered pairs in total,

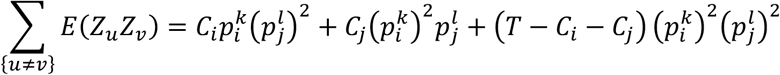

For *i* = *j* the two shared-cell cases coincide when *k* = *l* and are approximated by a single case otherwise.

The z-score is 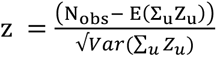, where *Nobs* is the observed whisper edge count.

The connexin analyses and the colon and skin scans use the independent-edge mode; the mouse brain ligand-receptor analysis uses the shared-cell mode; the two give nearly identical rankings (Spearman ρ = 0.999; Supp. Fig. 11).

#### Significance thresholds

Quadruplets are called at a fixed z-score threshold (z > 3 for connexin analyses, z > 4 for ligand-receptor analyses; under the normal approximation, *p* < 1.4 × 10⁻³ and *p* < 3.2 × 10⁻⁵) together with a minimum whisper network size of 30 edges. Because quadruplets share cells, cell types and genes, error control across the tested set is estimated empirically by the within-cell-type randomization control rather than from per-quadruplet p-values (see “Randomization and false positive rate estimation”).

### Stereoseq Data and Analysis

Mouse brain STEREO-seq data^27^ were obtained from the Spatial Omics Database^96^. After quality control filtering, the dataset comprised 50,140 cells and 18,527 genes, including 10 gap junction (connexin) genes: *Gjd2* (Cx36), *Gja5* (Cx40), *Gjc1* (Cx45), *Gja4* (Cx37), *Gjb2* (Cx26), *Gja1* (Cx43), *Gjb6* (Cx30), *Gjb1* (Cx32), *Gjc2* (Cx47), and *Gjc3* (Cx29). We used the authors’ annotated cell-type labels, encompassing 28 distinct cell types. Abbreviations include: CA3, cornu ammonis area 3; Mb, midbrain; EX, excitatory glutamatergic neuron; IN, GABAergic interneuron; DA, dopaminergic neuron; GN DG, granule cell of dentate gyrus; Astr, astrocyte; Micro, microglia; OPC, oligodendrocyte precursor cell; Oligo, oligodendrocyte; SMC, smooth muscle cell; Ery, erythrocyte; and Endo, endothelial cell.

Gene expression matrices were normalized and subsequently denoised using MAGIC imputation (t=2) to recover continuous expression profiles. To construct the *whisper network*, a gene was considered “expressed” in a given cell if its expression exceeded the 75th percentile of that gene’s expression across all cells. For gap-junction analysis, we evaluated all possible connexin– connexin combinations. Although several connexins are known to preferentially form specific homo-or heterotypic channels, experimental evidence for many pairings remains incomplete or context-dependent; thus, interaction patterns were inferred empirically from the data.

### Human colon and skin Xenium data

Xenium data for non-diseased human colon (Xenium Human Colon Gene Expression Panel, 425 genes) and human skin (Xenium Prime 5K Human Pan Tissue and Pathways Panel) were obtained from the 10x Genomics website and processed as above without imputation. The skin dataset is a primary dermal melanoma section; only the uninvolved epidermis and underlying dermis were analyzed (12,114 cells). The analyzed colon region comprised a 3.0 × 2.0 mm area (58,125 cells). Cell types were annotated from the panel-provided marker gene sets and graph-based clusters. In colon, goblet cells were resolved by subclustering the epithelial clusters, and stromal classes (fibroblast, smooth muscle, pericyte, endothelial) were assigned from canonical markers. In skin, keratinocytes were separated into basal (TP63, COL17A1, ITGB4, ITGA6, LAMB3) and suprabasal (DMKN, CDSN, DSG1) cells by marker score, and the suprabasal population was divided into three layers by tertiles of distance to the nearest basal cell; keratinocyte-scored cells more than 250 µm from the basal layer were excluded. A positional zonation score was computed for every colon cell as the mean of surface-marker (CA1, CA2, AQP8, SLC26A3, GUCA2A, GUCA2B, MS4A12, SELENBP1) minus crypt-base-marker (LGR5, OLFM4, ASCL2, SOX9, MKI67, PCLAF) expression over its ten nearest epithelial cells. All CellChatDB interactions (human, cell-cell contact) with both partners being single genes present on the panel were tested (colon, 9 pairs; skin, 145 pairs) with default settings (75th-percentile binarization, k = 5). Quadruplets with z > 3 and whisper network size > 30 are reported in Supp. Data 11 and 12, together with the fraction of cells of each cell type in which each gene is detected; quadruplets in which a gene is detected in under 5% of a cell type most likely reflect transcript misassignment at cell boundaries and are flagged.

### Xenium mouse brain data (wild-type and TgCRND8)

Xenium In Situ data for coronal sections of wild-type and TgCRND8 (Alzheimer’s disease model) mouse brain were obtained from 10x Genomics, one section per condition, at the terminal time point of the published time course (age >13 months), at which amyloid pathology is well established. Data were processed as above without imputation. Of the panel genes, three annotated gap junction genes (Gja1, Gjb2, Gjc3) were used as signaling genes. Cross-sample integration and cell-type label transfer are described under “Comparison of conditions (WT vs TgCRND8)” below.

### Ligand–receptor database

We used the CellChat v2^22^ ligand–receptor compendium, focusing specifically on pairs annotated as cell–cell contact interactions. After intersecting this database with the gene set retained from the STEREO-seq dataset, 367 ligand–receptor pairs were included for downstream analyses.

### Randomization and False Positive Rate estimation

We performed randomization by permuting cell IDs within each cell type. This procedure preserves the marginal spatial distribution and gene expression profiles specific to each cell type. False positive rate (FPR) is obtained by comparing the number of detected pairs obtained on randomized data with number of detected pairs on real data.

### Sensitivity to method parameters

To assess the dependence of results on preprocessing and graph parameters, we varied one setting at a time around the default configuration (MAGIC t = 2, 75th-percentile binarization, k = 5): imputation (none, MAGIC t = 1, 2, 3), binarization percentile (70, 75, 80) and neighborhood size (k = 5, 7, 9). For each of the eight configurations we reran the full connexin analysis (100 gene pairs, 28 cell types) on the STEREO-seq dataset and on its within-cell-type randomized counterpart, and recorded the number of significant quadruplets (z > 3, whisper network size > 30), the empirical false-positive rate, the fraction of default-configuration findings retained, and the Spearman correlation of quadruplet z-scores with the default (Supp. Fig. 8). Imputation is applied only to the sequencing-based STEREO-seq data, where transcript dropout otherwise leaves too few connexin-expressing cells to form whisper networks; Xenium datasets are analyzed without imputation. MAGIC’s diffusion parameter was set to t = 2 rather than the default t = 3 to limit over-smoothing, since smoothing-induced spatial coherence is preserved by within-cell-type randomization and therefore cannot be detected by the randomized control.

### Robustness to annotation errors

To assess sensitivity to cell-type annotation errors, we relabeled 1, 2, 5 or 10% of cells in the STEREO-seq dataset and reran the connexin analysis with default settings. Two error modes were used: “random”, in which a relabeled cell receives a label drawn uniformly from the other cell types, and “neighbor”, in which it receives the label of a randomly chosen spatial neighbor of a different type, mimicking boundary and segmentation-driven mis-annotation. Each setting was repeated with three random seeds. We report the fraction of quadruplets significant under the original annotation that remain significant, and the Spearman correlation of z-scores with the original annotation (Supp. Fig. 9).

### Benchmarking

CellChat v2 and COMMOT were run on the full 50,140-cell STEREO-seq dataset with a custom database of the 100 ordered connexin pairs used by CellWHISPER, annotated as cell-cell contact interactions. SpaTalk did not complete on the full dataset even with 1.5 TB of memory, and COMMOT’s joint optimization did not complete for the full ligand-receptor compendium; the comparison in Fig. 1d therefore uses a 5K-cell hippocampal region (Supp. Fig. 5) for all three tools, randomized by permuting cell IDs within each cell type.

All comparison methods were downloaded from their official repositories and executed using recommended parameters. For CellChat v2, we used the computeCommunProb function with default parameters (type = "triMean", trim = 0.1, interaction.range = 250, contact.knn.k = 5), followed by filterCommunication with a minimum of 20 cells per cluster. For COMMOT, we applied commot.tl.cluster_communication to compute cell–cell communication for each ligand–receptor pair. For SpaTalk, we used the default parameters.

### Prioritization of reference interactions (Supp. Fig. 10)

As reference we used the five interactions with independent support in this study: the three Cx43 interfaces confirmed by immunostaining (Fig. 3) and the two oligodendrocyte gap junctions with literature support (Fig. 2c). For each method, quadruplets were ranked by its own score (z-score for CellWHISPER; p-value for CellChat v2 and COMMOT, with tied quadruplets counted as a block), and we recorded how many quadruplets a user would need to screen to recover all five. CellChat v2 and COMMOT assign their minimum p-value to 12,358 and 9,680 quadruplets respectively, reporting only 5 distinct p-values across their significant output, against 2,298 distinct z-scores for CellWHISPER. A block of ties this large indicates a null that is violated broadly: both methods test against a null that does not preserve cell-type-specific expression and spatial organization, so cell-type co-localization alone pushes many quadruplets past every permuted value (Fig. 1d). More permutations would break the ties, but the ranking would still reflect departure from this null rather than contact-specific co-expression.

### Connexin mRNA-protein concordance

CellWHISPER relies on cellular mRNA abundance as a proxy for membrane-bound connexin proteins. To evaluate this assumption, we performed targeted spatial profiling of Cx43 protein and its corresponding mRNA (GJA1) at single-molecule resolution in an induced pluripotent stem cell (iPSC) model, where gap junction communication is critical for spatial patterning.

We observed a significant correlation between GJA1 mRNA expression and average cytosolic Cx43 protein intensity at the single-cell level (Spearman’s R = 0.47, p = 5 × 10⁻²⁰; Supplementary Fig. 1). Consistently, whisper networks inferred from GJA1 mRNA showed strong overlap with those inferred from Cx43 protein measurements (Fig. 1e, Supp. Fig. 1), supporting the use of mRNA as a proxy for connexin-mediated communication in this context.

### Latent Variable Model (LVM)

LVM stipulates that the binary variable *I (ct_i_*, *ct*_j_, *g_k_*, *g_l_*) indicating whether the CT-SG quadruplet *ct_i_*, *ct*_j_, *g_k_*, *g_l_* has statistical support in the data follows a Bernoulli distribution:

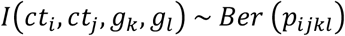

where *u_i_*, *u*_j_ denote to-be-learnt low-dimensional representations (embeddings) for CTs *ct_i_*, *ct*_j_and *v_k_*, *v_l_* denote embeddings for SGs *g_k_*, *g_l_*, respectively, all embeddings being in a common latent space. The dot product *u_i_*. *v_k_* captures the general propensity of *ct_i_* for participating in CCC using *g_k_*. Thus, the model posits that the quadruplet *ct_i_*, *ct*_j_, *g_k_*, *g_l_* is more likely if there is a higher tendency for *ct_i_* communicating via *g_k_* and *ct*_j_ communicating via *g_l_*.

Using this formulation CellWHISPER learn embeddings that enable quantification of (1) cell type-signaling gene preferences (*u_i_*. *v_k_*), (2) cell type similarities (*u_i_*. *u*_j_), i.e., which pairs of CTs have similar communication mechanisms, and (3) signaling gene similarities (*v_k_*. *v_l_*), i.e., which SGs are used by the same cell types. It uses cosine similarity rather than dot products between embeddings to emphasize specificity and mitigate abundance bias inherent in dot product.

In the null model, *p_i_*_j*kl*_ is assumed to be constant for all indices *i*, *j*, *k*, *l*. In contrast, the alternate model factorizes *p_i_*_j*kl*_ into two terms:

- the propensity of cell type *i* to communicate via signaling gene *k*, and
- the propensity of cell type *j* to communicate via signaling gene *l*

Formally, this is expressed as:

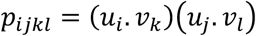

where *u_i_* and *u*_j_ are low dimensional embeddings for cell types *i*, *j*, and *v_k_*, *v_l_* are embeddings for signaling genes *k* and *l*. All embeddings lie in a shared latent space, and we enforce the constraints:

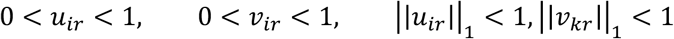

This formulation is invariant under simultaneous permutation of both CT and SG pairs, i.e.

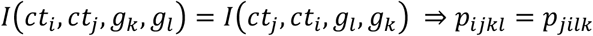

We optimize the model using constrained log-likelihood maximization, implemented via `scipy.optimize` using the SLSQP method. The constraint is posed using the L1 norm of the embeddings [|*u_i_*|[_1_ = *Σ_d_u_id_* (entries being positive), which keeps the constraints linear and ensures probabilities 0 < *p_i_*_j*kl*_ < 1.

To confirm identifiability and recovery, we simulated Gap Junction Communication results with the same orders as the STEREO-seq analysis (29 cell types, 10 connexins, d=3). Ground-truth embeddings were drawn from numpy.random.rand with unit norm cell type and signaling gene embeddings. The optimizer recovered CT–CT similarity, SG–SG similarity, and CT–SG preference structure with high agreement (Fig. 5b).

For ligand–receptor analysis, only a subset of ligand–receptor pairs are biologically plausible. To ensure model learnability, we explicitly set the probability *p_i_*_j*kl*_ = 0 for incompatible ligand– receptor combinations. For computational feasibility, we randomly subsample a small number of such incompatible pairs (2000 in our experiments) to include as negative examples during optimization. This approach enables efficient training while maintaining biological specificity in the learned embeddings.

### Comparison of conditions (WT vs TgCRND8)

We compared wild-type and TgCRND8 mouse brains, one Xenium section each, at the terminal time point at which amyloid pathology is well established. Wild-type (WT) and TgCRND8 (AD) Xenium datasets were jointly integrated using residualPCA to achieve conservative alignment across conditions while preserving biological differences. A joint embedding was constructed and cell-type clustering was performed on the integrated space yielding 25 clusters.

Label transfer from reference. Cell-type labels were transferred from the annotated STEREO-seq reference (analyzed above). For this step, the WT, AD, and STEREO-seq datasets were co-embedded with reciprocal PCA^97^. For each query cell (WT or AD), we identified its k = 7 nearest neighbors in the reference and recorded their labels. For each cluster in the joint WT+AD embedding, neighbor votes were aggregated across all member cells and normalized by the total number of votes in that cluster to yield a label-probability vector. The cluster label was assigned as the cell type with the maximum normalized vote. Clusters containing multiple subpopulations of the same broad class (e.g., oligodendrocytes) were further subdivided and relabeled as subtypes (e.g., Olig-1, Olig-2).

#### Per-condition quantification

Using these harmonized annotations, CellWHISPER quantified CT– SG quadruplets separately in WT and AD datasets. Quadruplets were retained if they satisfied significance thresholds (*z*-score > 4 and whisper-network size > 30). This procedure yielded 60 high-confidence quadruplets across the two conditions. A quadruplet was classified as shared if both WT and AD satisfied these thresholds, condition-specific if one condition satisfied the thresholds while the other did not (*z*-score < 3 or whisper-network size < 20), and indeterminate otherwise. Condition-specific here means detected in one condition only, not a tested difference between conditions. Since only a single section is available per condition, these are descriptive comparisons between two samples rather than tests of condition-specific communication.

### Experimental details for Cx43 protein and its mRNA correlation iPSC culture

iPSCs were obtained and cultured according to Culture and Freezing Methods for WTC Derived AICS hiPSC Lines: https://www.coriell.org/0/PDF/Allen/iPSC/AICS_SOP_WTC_CellCulture.pdf. Coverslips (22 × 22 mm) were placed flat in sterile 6-well plates and coated with growth factor– reduced, phenol-red–free Matrigel (Corning, cat. no. 356231) at a final protein concentration of 0.337 mg/mL. Briefly, Matrigel aliquots thawed at 4 °C were diluted on ice in cold DMEM/F12 to 1.5 mL per well, mixed thoroughly with pre-chilled pipettes to ensure homogeneity, and immediately dispensed to fully cover each coverslip. Plates were incubated at room temperature for 1 h, after which excess coating solution was aspirated and wells were gently rinsed with room-temperature mTeSR1 medium supplemented with 1 µM Y-27632 (ROCK inhibitor) before cell seeding. Coated coverslips were used for experiments within 6 h or, if needed, sealed with Parafilm and stored at 4 °C for up to 2 weeks. Once cells were confluent, they were fixed in 4% formaldehyde for staining.

### iPSC RNA and protein multiplexing

Fixed cells on coverslip with permeabilized with 70% EtOH for 30 mins at -20 C. Cycle 1 RNA was labeled using Molecular Instruments hybridized chain reaction RNA fluorescence in situ hybridzation (HCR RNA FISH) v3.0 probes: Human Cx43 B3 555, Human MALAT1 B2 647. All probes were amplified 75 mins at room temperature. Fluorescence images were acquired on adapted Nikon Ti 2000-U microscope with appropriate filters. Then, samples were blocked and incubated with Cx43 Ab (Invitrogen 71-0700, 1;150) overnight at 4 C followed by secondary antibody staining (anti-rabbit 488) for 1 hr at room temperature. This Cycle 2 Cx43 Ab was imaged again. DAPI counterstain was used to register the images from both cycles. Images were segmented with Cellpose and quantified for downstream analysis.

### Immunofluorescence staining

Mouse brain sections were fixed in 4% paraformaldehyde (PFA) and sectioned using Slides were deparaffinized and rehydrated by subsequent rinses in Xylene, 100% EtOH, 95% EtOH, 80% EtOH, 70% EtOH and deionized water (diH_2_O). Then we performed antigen retrieval by placing slides in a Coplin jar with 10% sodium citrate buffer in 1x phosphate-buffered saline (PBS). Coplin jars were placed in a pressure cooker with 300 mL of tap water and set on high pressure for 15 minutes. The slides were let to cool for 20 minutes and then washed with diH_2_O. They were then blocked using 50 mg/mL bovine serum albumin (BSA) in 1x PBS. They were then washed with .1% Tween 20 in 1x PBS (PBS-T). Primary antibodies were diluted based on manufacturers’ recommendation in 5% BSA and left to incubate on slides at 4°C, overnight. A list of antibodies and their dilutions are below. Slides were washed the next day with PBS-T. For unconjugated primary antibodies, slides were washed with three times with PBS-T for 2 minutes and incubated in 1:5000 <u>secondary antibody</u> in PBS for 1 hour at room temperature. Slides were washed were washed three times with PBS-T and counterstained by a 1:500 Dapi in 1x PBS solution for 10 minutes. Slides were washed with PBS-T for 2 minutes and nuclear staining was done using 1:2000 DAPI (Thermo Scientific, Ref:62248). Slides were mounted using 10% glycerol in 1x PBS. Coverslips were placed over top of the slides, and the edges were coated with nail polish to prevent dehydration. Slides were imaged using the Keyence BZ-X800 fluorescence microscope.

### Image preprocessing and quantification of colocalization

Images of cells in tissue all underwent the following preprocessing before further analysis. Each image contains multiple channels. The maximum intensity projection of each channel was then generated. The projected image was the final image used for cellular counting analysis. For the DAPI channel, the maximum intensity projection was generated from the raw image. For visualization consistency, brightness and contrast of each fluorescent channel were adjusted uniformly across all images using the “Brightness/Contrast” tool without altering the underlying pixel intensity values. To measure intercellular colocalization, the straight-line tool was used to draw a line between the centers of identified cells of interest (e.g., nuclei or marker-specific regions). The length of each line was recorded using the “Measure” function, which reports distances in calibrated units (µm) based on the microscope’s pixel-to-micrometer conversion. At least 629 cells per field were analyzed, and distances were exported for downstream statistical analysis. Previous steps were conducted in ImageJ/Fiji (National Institutes of Health). Statistical significance was calculated using a regular two-way ANOVA in *BioRender Graph*. Statistical levels are provided as follows: ns for 0.05 < p ≤ 1, * for 0.01 < p ≤ 0.05, ** for 0.001 < p ≤ 0.01, *** for 0.0001 < p < 0.001, and **** for p ≤ 0.0001.

### Availability of data and materials

Mouse brain STEREO-seq data were obtained from the Spatial Omics Database (https://db.cngb.org/stomics/). Xenium TgCRND8 Alzheimer’s disease mouse model data were obtained from 10x Genomics. iPSC-derived and mouse brain immunostaining data will be made available upon publication. The source code for CellWHISPER is publicly available at https://github.com/anurendra/CellWHISPER under an open-source license.

## Acknowledgements

This work was supported by the National Institutes of Health (R35GM131819 to S.S.; R35GM151028 to A.F.C.) and the National Science Foundation CAREER award (2338935 to A.F.C.). A.F.C. also acknowledges support from the Burroughs Wellcome Fund Career Award at the Scientific Interface, the Bernie Marcus Early-Career Professorship, and start-up funds from Georgia Institute of Technology and Emory University. The authors thank Dave Zhao for helpful discussions and feedback on the manuscript.

## Authors’ contributions

AK conceived the conceptual formulation and validation framework. AK and SS developed the mathematical formulation. AK implemented and tested the method. AK prepared all figures except figure 3. BA contributed to experimental analyses. FR, NZ, and AC designed and performed experimental validation in iPSC and mouse brain models. FR prepared figure 3. AK and SS wrote the manuscript. All authors read and approved the final manuscript.

## Declaration of generative AI and AI-assisted technologies in the manuscript preparation process

During the preparation of this work the authors used Claude (Anthropic) to improve the readability and formatting of text. After using this tool, the authors reviewed and edited the content as needed and take full responsibility for the content of the publication.

