## Supplementary Figures for "CellWHISPER disentangles direct cell–cell communication from structural proximity"

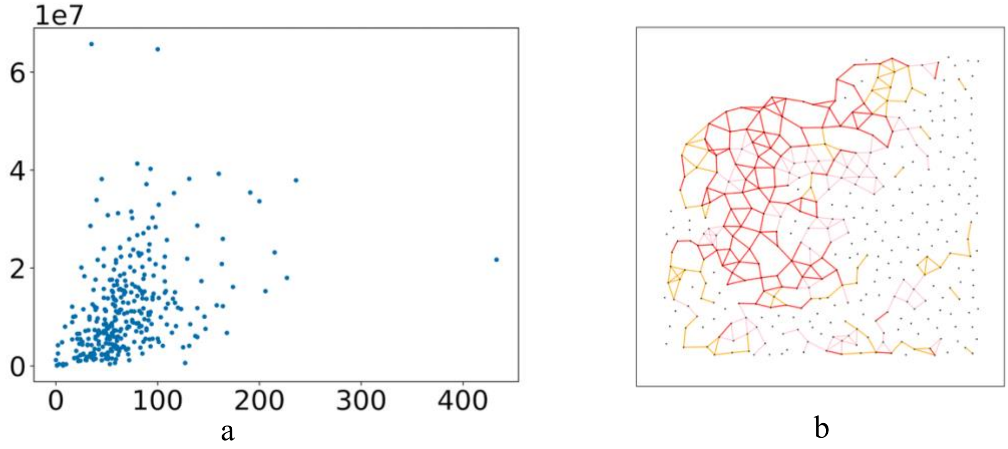


**Supplementary Figure 1:** (a) Gap junction gene-protein concordance: Gja1 mRNA expression correlates with cytosolic concentration of its encoded connexin protein Cx43 (Spearman correlation = 0.47), as measured in our induced pluripotent stem cell (iPSC)- derived cultures using targeted spatial profiling. (b)Whisper network analysis on an in-house iPSC cell line dataset using cytosolic Cx43 protein and GJA1 mRNA. Pink edges represent whisper edges exclusive to cytosolic Cx43 protein, orange edges indicate those specific to GJA1 mRNA, and red edges highlight shared connections between both networks. A 50th percentile threshold was applied.

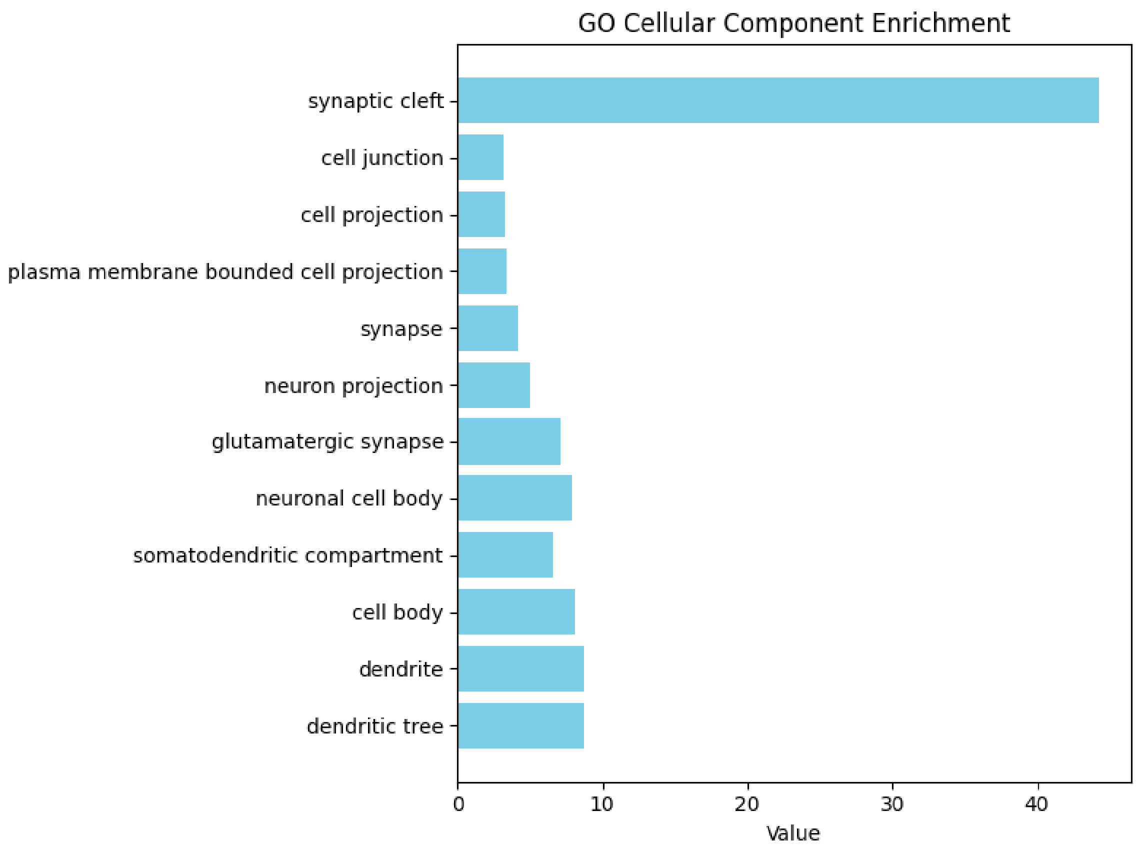


**Supplementary Figure 2**. Fold enrichment of randomly selected genes with higher edge counts across GO cellular component categories shows enrichment of direct cell junction and ECM related terms. Only terms with $p < 0.005$ are shown.

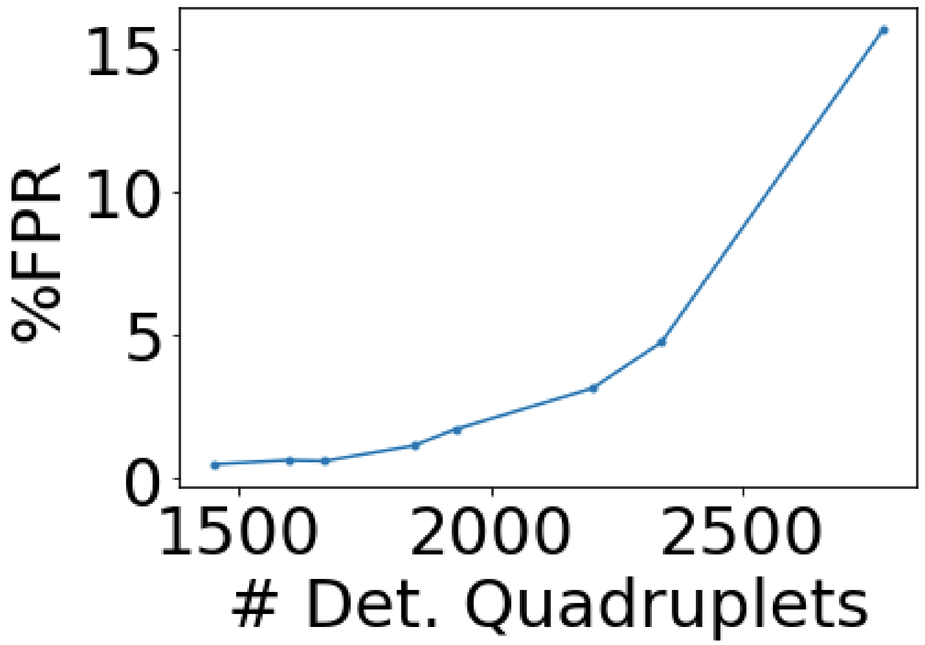


**Supplementary Figure 3:** Empirical false positive rate (%FPR, y-axis) plotted against the number of detected CT–SG quadruplets (x-axis), as the significance threshold is progressively relaxed on mouse brain STEREO-seq data (~50,000 cells, 28 cell types, 10 connexin genes). FPR is defined as the number of significant quadruplets detected on within-cell-type randomized data divided by the number detected on real data. At the operating threshold used in this study (z-score > 3, whisper network size > 30, ~1,726 quadruplets), FPR remains below 1%. FPR increases sharply as thresholds are relaxed beyond ~2,000 quadruplets, indicating that the stringent operating point provides well-controlled inference while retaining biologically meaningful signal.

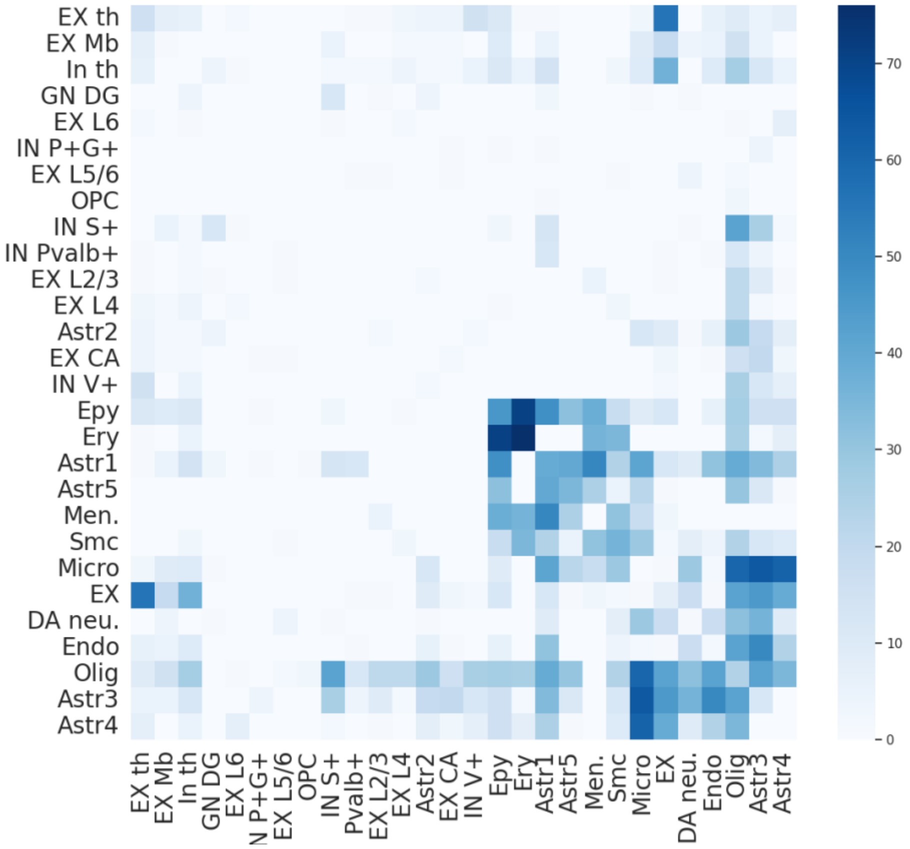


**Supplementary Figure 4:** Cell type similarity derived from aggregated cell–cell communication across all gap junction pairs does not reveal a modular structure. Cell types were ordered by performing hierarchical clustering on the aggregated matrix.

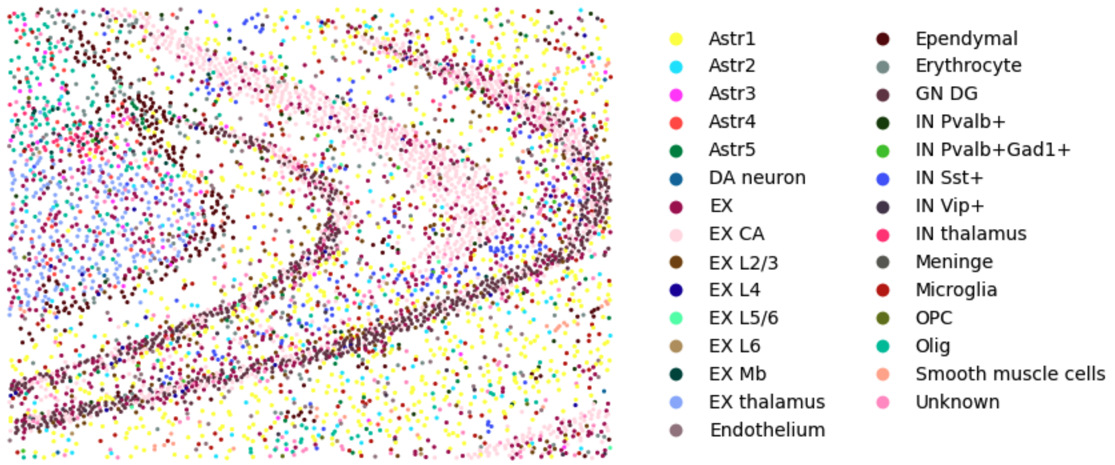


**Supplementary Figure 5:** Spatial plot of ~5,000 hippocampal cells from the mouse brain STEREO-seq dataset used in benchmarking experiment (Fig 1d).

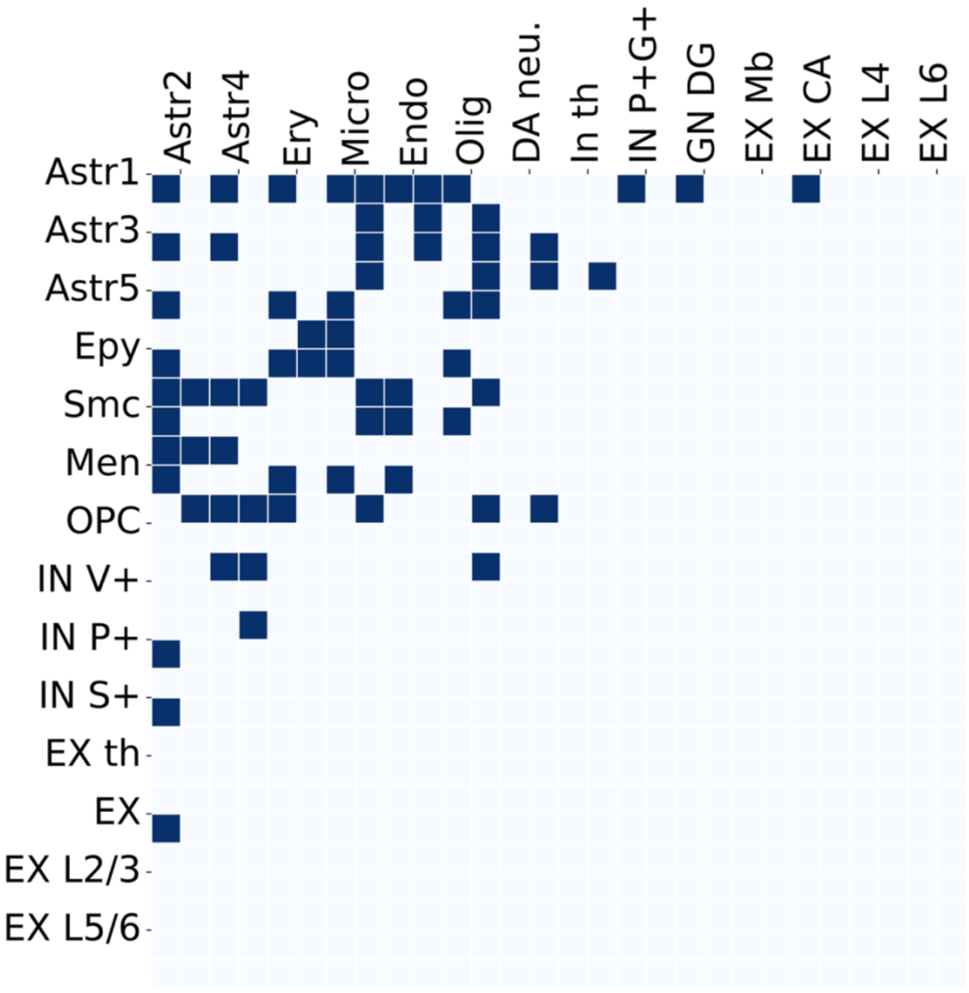


**Supplementary Figure 6:** Full map of cell type pairs predicted to communicate via homotypic Cx43–Cx43 gap junctions. Each filled square indicates a significant CT–SG quadruplet (z-score > 3, whisper network size > 30) involving homotypic Cx43 (GJA1) on both sides of the junction, in the STEREO-seq mouse brain dataset. Communication is predominantly glial: astrocyte subtypes (Astr1, Astr3, Astr5) show the broadest connectivity, communicating with microglia, endothelial cells, oligodendrocytes, ependymal cells, and smooth muscle cells. Neuronal Cx43 communication is sparse, consistent with the known predominance of Cx43 in glial rather than neuronal coupling. This figure complements Fig. 2g, which shows the subset of glial interactions highlighted in the main analysis.

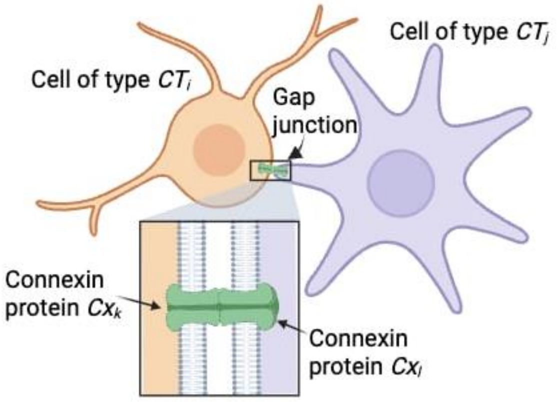


**Supplementary Figure 7: Schematic of Gap junction communication:** Connexin proteins dock at the membranes of adjacent cells, potentially of different cell types ($CT_i$, $CT_j$) to form intercellular channels called “gap junctions”, enabling direct communication via transport of small molecules such as ions, metabolites, and second messengers. The connexin proteins ($Cx_k$, $Cx_l$) may be homotypic or heterotypic.

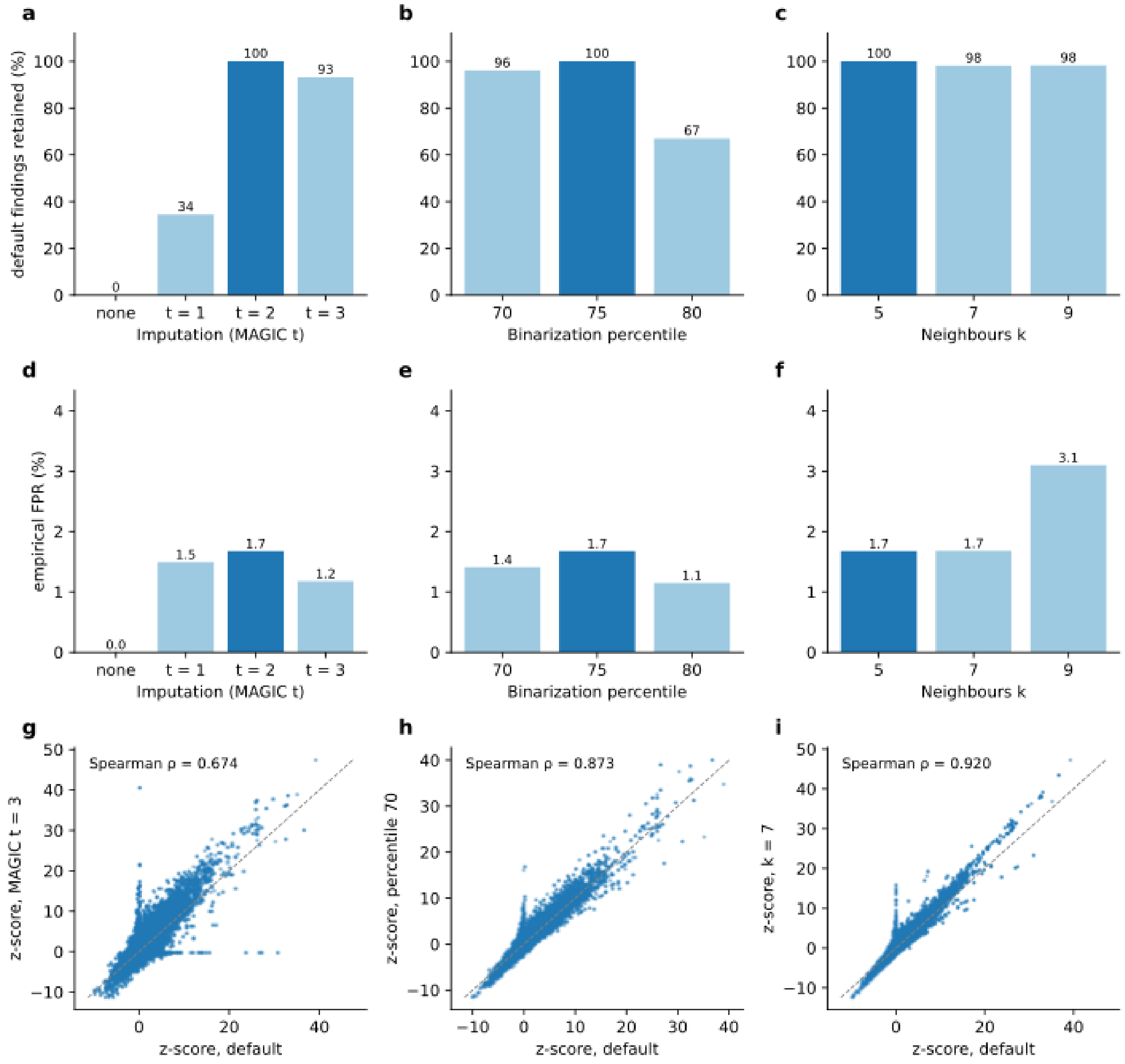


**Supplementary Figure 8: Sensitivity of CellWHISPER to hyperparameters.** All 100 connexin pairs were tested on the STEREO-seq mouse brain section, varying one parameter at a time around the default (MAGIC t = 2, 75th-percentile binarization, k = 5; dark bars). (a-c) Fraction of quadruplets significant at default (z > 3, whisper network size > 30) that remain significant in each configuration. (d-f) Empirical false-positive rate, the number of significant quadruplets on within-cell-type randomized data divided by the number on real data. (g-i) Quadruplet z-scores in the default configuration against MAGIC t = 3, percentile 70 and k = 7, with Spearman correlation.

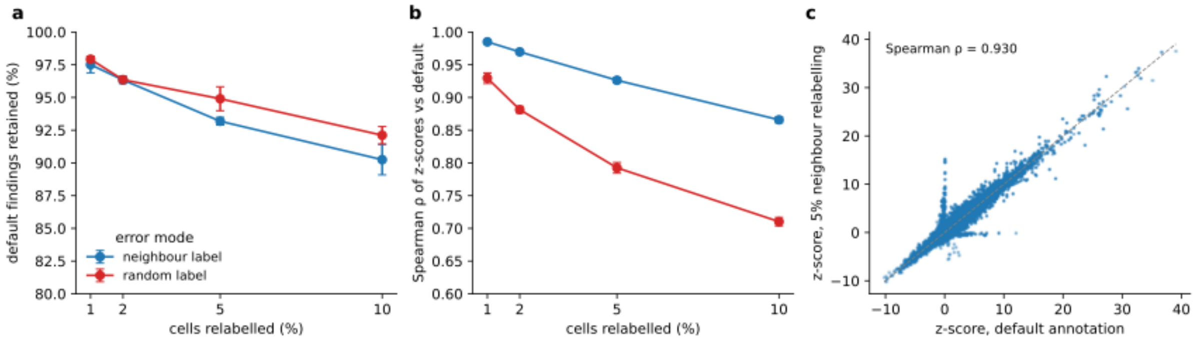


**Supplementary Figure 9: Robustness to cell-type annotation errors.** A fraction of cells in the STEREO-seq brain section was relabelled, either at random or with the label of a spatial neighbour, and all 100 connexin pairs were rerun (3 seeds; mean ± s.d.). (a) Fraction of quadruplets significant with the original annotation ($z > 3$, whisper network size > 30) that remain significant. (b) Spearman correlation of quadruplet z-scores with the original annotation. (c) Quadruplet z-scores with the original annotation against 5% neighbour relabelling (one seed).

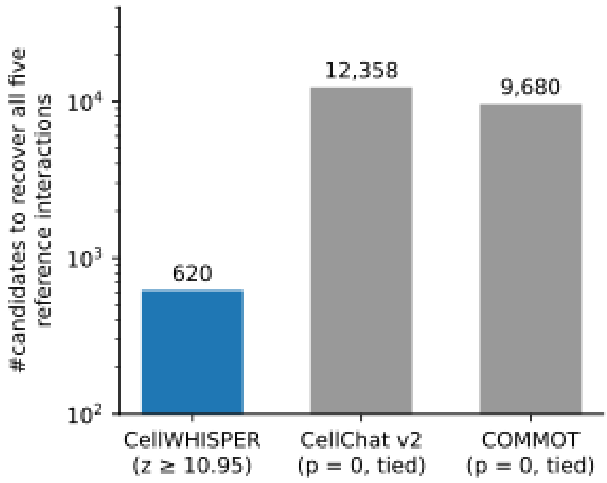


**Supplementary Figure 10: Prioritization of reference interactions.** Number of top-ranked quadruplets a user must screen to recover five reference interactions: the three Cx43 interfaces supported by immunostaining (Figure 3; astrocyte–astrocyte, astrocyte–endothelium, astrocyte–microglia, Gja1-Gja1) and two established oligodendrocyte gap junctions (Figure 2c; Olig–Olig Gjb1-Gjb1 and Gjc2-Gjb1). Full 50K-cell STEREO-seq dataset, 100 connexin pairs. CellWHISPER ranks by z-score (all five recovered at z ≥ 10.95, 620 quadruplets); CellChat v2 and COMMOT rank by p-value, and all five fall within the set tied at the minimum p-value (12,358 and 9680 quadruplets).

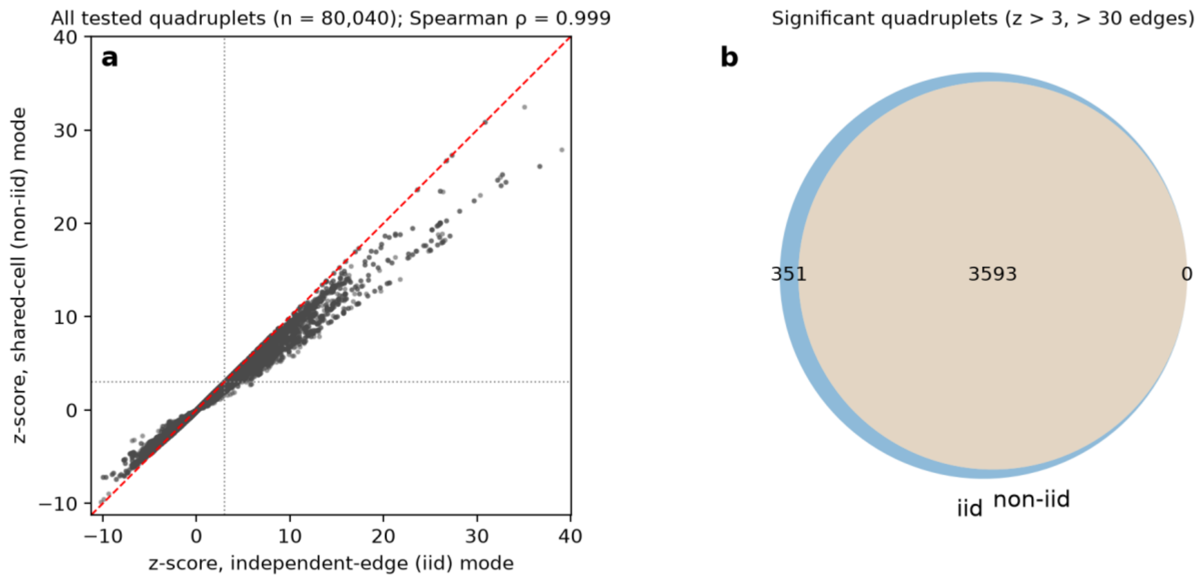


**Supplementary Figure 11: Independent-edge and shared-cell variance give near-identical z-scores.** (a) Quadruplet z-scores for all 100 connexin pairs on the STEREO-seq brain section (80,040 quadruplets) computed with the independent-edge (iid) variance against the shared-cell (non-iid) variance. Accounting for shared cells increases the variance, so non-iid z-scores are systematically slightly smaller, while the ranking is preserved (Spearman ρ = 0.999). (b) Significant quadruplets (z > 3, whisper network size > 30) under the two modes: the non-iid set (3,593) is a subset of the iid set (3,944).

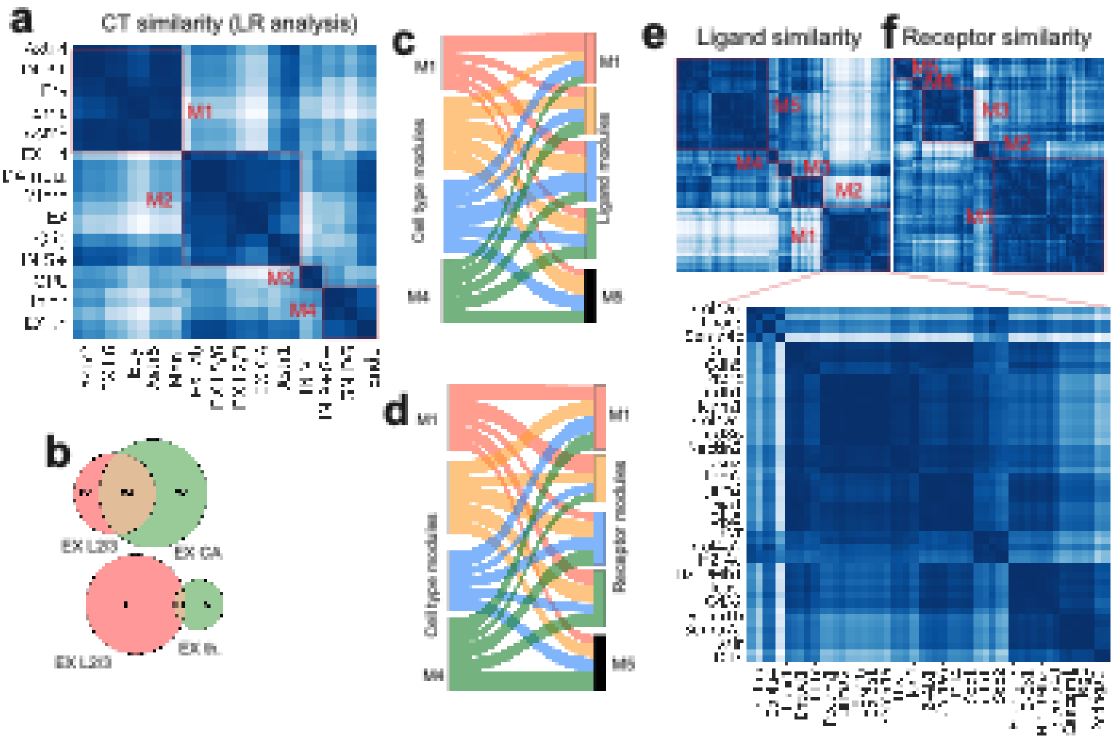


**Supplementary Figure 12: LVM analysis of ligand–receptor communication in the Stereo-seq mouse brain**. (a) Cell-type similarity matrix for ligand–receptor communication. Cosine similarity of latent cell-type embeddings derived from CellWHISPER applied to ligand–receptor interactions (Stereo-seq mouse brain dataset). Four dominant cell-type modules emerge. Excitatory neurons in cortical layers and hippocampal CA region cluster together (M2), whereas excitatory neurons in thalamus is in module M4. (b) Communication overlap between excitatory neurons. Quantification of overlap in ligand–receptor communication patterns (shared CT–ligand–receptor triplets) shows that excitatory neurons in cortical layers 2/3 are more similar to CA excitatory neurons than to thalamic excitatory neurons, consistent with known laminar and neuromodulatory properties of neocortical and hippocampal neurons. (c–d) Cell type–ligand and cell type–receptor module specificity. Sankey plots showing preferential associations between cell-type modules and ligand modules (c), and between cell-type modules and receptor modules (d). Links were derived by averaging CT–SG preference scores within each module. (e) Ligand similarity map. Cosine similarity of ligand embeddings reveals five dominant modules (Supp. Data 9). Module M1 consists mainly of genes related to cell adhesion (e.g., PECAM1, CDH1/5, ICAM1), extracellular matrix components (COL4A1–6, LAMA2), and immune modulators (H2-Aa, H2-Eb1, H2-Ab1, CD52, CD55, CD86, PTPRC). Another ligand module (M5, Supp. Data 9) is enriched for axon guidance and synaptic signaling genes, including protocadherins, ephrins, neurexins, semaphorins, Lrrc and Flrt family. (f) Receptor similarity map. Cosine similarity of receptor embeddings identifies five receptor modules (Supp. Data 9).
